# Citrate-Based Hydrogels Promote Closure of Infected Diabetic Wounds

**DOI:** 10.64898/2026.11.06.764741

**Authors:** Maria Mendez-Santos, Marilene van den Berg, Ophelia Tong, Caralyn Collins, Ziyou Ren, Guillermo A. Ameer

## Abstract

Diabetic foot ulcers (DFUs) are a significant threat to the quality of life for individuals with diabetes, affecting approximately 1.6 million patients in the US, as of 2023. When these wounds do not heal and become chronic wounds, they can become infected and increase the risk of digit and/or lower limb amputation. Therefore, there is a critical need for therapies that promote wound closure in the presence and absence of infection. We have previously reported the potential benefits of the thermoresponsive citrate-based macromolecule poly(polyethylene glycol citrate-co-N-isopropylacrylamide) (PPCN) in diabetic rodent and swine models of non-infected full thickness dermal wounds. In this study, we compared four PPCN formulations (PPCN, PPCN-CMOF, A5G81-PPCN, and A5G81-PPCN-CMOF), which are functionalized with the laminin-derived cell adhesion peptide A5G81 and/or contain entrapped copper metal-organic framework nanoparticles (CMOF) designed to release Cu²+ slowly, in diabetic mice with wounds infected with *S. aureus*. In infected wounds, all four formulations significantly improved cumulative closure relative to saline (adjusted p = 0.002–0.021). In non-infected wounds, only PPCN-CMOF significantly improved cumulative closure relative to saline (p = 0.046), although all four formulations reached ≥95% closure earlier than saline (adjusted p = 0.009–0.045). *In vitro*, the CMOF-containing formulations produced the highest bacterial killing (90% for A5G81-PPCN-CMOF and 84% for PPCN-CMOF), whereas A5G81-PPCN and PPCN reduced bacterial colonies by 76% and 59%. PPCN modification with A5G81 and/or CMOF increased endothelial mesh formation *in vitro*. *In vivo*, PPCN-based formulations increased granulation tissue thickness and, depending on formulation and infection status, vascular marker expression. These results suggest that PPCN and PPCN formulations containing CMOF and/or A5G81 are a potentially viable technology that can be added to the wound care toolbox to accelerate healing of diabetic foot ulcers in the presence of infection. These findings support further investigation of PPCN as an instructive pro-regenerative biomaterial in a preclinical large animal model.

**Graphical abstract:** 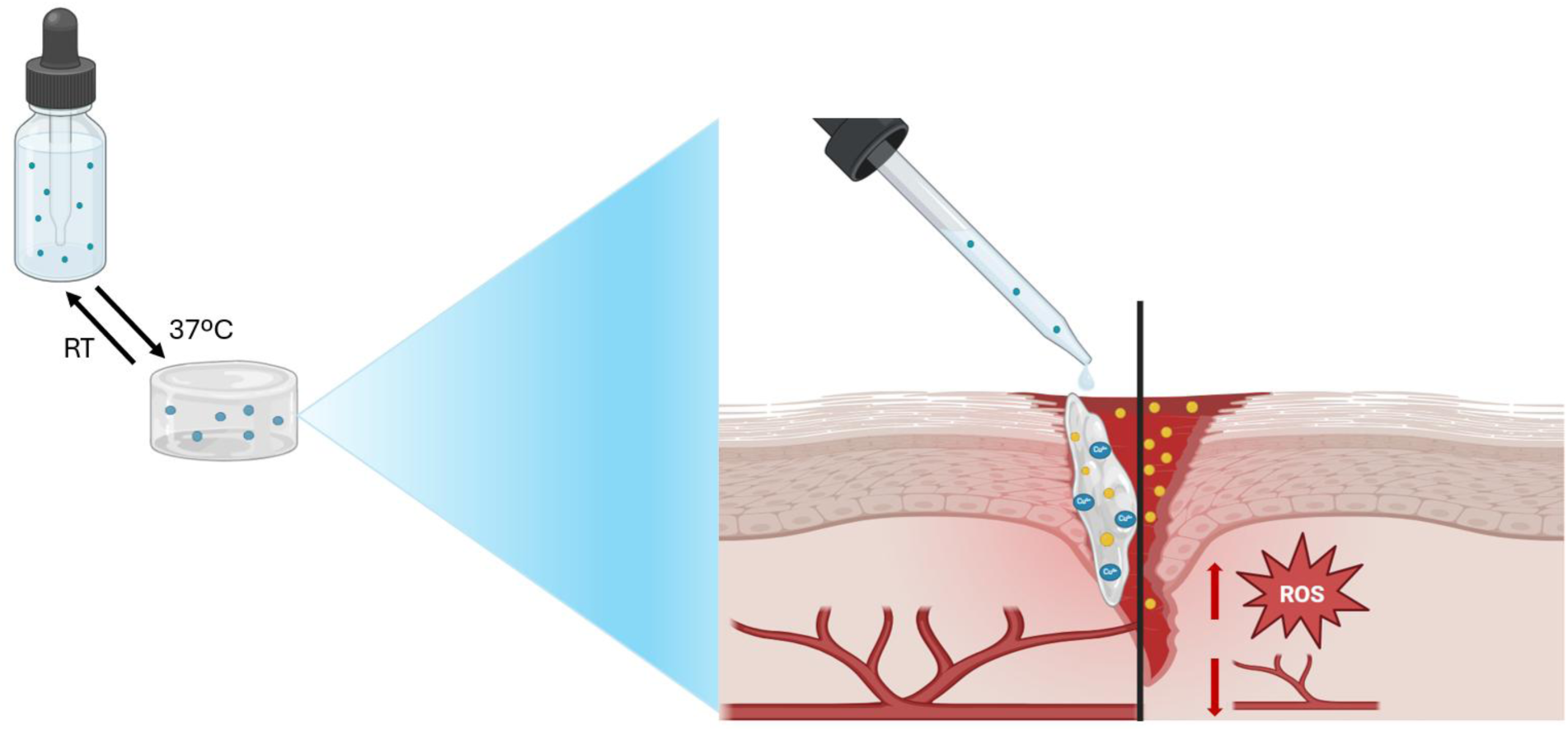

**Statement of Significance:** DFUs represent one of the most devastating complications of diabetes, as they are often infected, and may result in lower-limb amputation. A major barrier to effective healing of DFUs is the multifaceted pathophysiology, characterized by impaired vasculaturization, uncontrolled inflammation and oxidative stress, and peripheral neuropathy. This study set out to address these challenges by evaluating a citrate-based thermoresponsive hydrogel (PPCN) functionalized with the peptide A5G81 and/or copper metal-organic framework nanoparticles (CMOF), designed to be antioxidative, antibacterial, vasculoinductive, and dermoconductive. We show that PPCN-based hydrogels can improve closure of infected wounds in diabetic mice.

## Introduction

Diabetes affects more than 40 million people in the US^1^ and costs an estimated $413 billion annually, including more than $100 billion in indirect costs from disability-related job loss, reduced on-the-job productivity, and premature death^2^. Diabetic foot ulcers (DFUs) are among its most debilitating complications: about 1.6 million people in the US develop one each year, roughly half of these ulcers become infected, and most lower-extremity amputations in people with diabetes are preceded by an ulcer^3^. Beyond the risk of limb loss and death, DFUs markedly reduce physical function, vitality, and overall quality of life^4^, imposing a heavy personal and societal burden.

DFUs begin with neuropathy-related loss of protective sensation, which permits repetitive, unnoticed trauma^3^, and peripheral artery disease frequently coexists^5^. Once the wound forms, hyperglycemia-driven oxidative stress and endothelial dysfunction impair angiogenesis, cell migration, and granulation tissue formation, and hold the wound in a prolonged inflammatory state^6^. The open wound also provides a favorable niche for bacterial colonization, and diabetes-associated immune dysfunction limits host defense^6,7^, making DFUs prone to persistent and drug-resistant infection, especially by *Staphylococcus aureus*^8^. Infection in turn prolongs inflammation, damages tissue, and raises the risk of amputation and death^3^.

Research strategies to address this problem fall into three broad categories. The first is antimicrobial control of the wound, including antiseptics, silver-or iodine-containing dressings, antibiotic-or antimicrobial peptide-loaded materials, and metal-based nanoparticles^9,10^. The second is biologic stimulation of repair, using recombinant growth factors and cytokines, cell-based therapies, and skin substitutes^11^. The third is multifunctional biomaterials, mainly hydrogels, that combine antibacterial activity with antioxidant, pro-angiogenic, or pro-migratory cues, often through bioactive peptides, metal ions, or metal-organic frameworks^12^.

Despite this activity, few of these approaches have become clinically useful, and the certainty of evidence for interventions that enhance DFU healing remains poor overall^13^. Antimicrobial approaches reduce bacterial burden but do not regenerate tissue, and some raise concerns about cytotoxicity and microbial resistance^14^. Biologics are limited by short half-lives, instability, rapid diffusion, and uncontrolled signaling^12^. Multifunctional biomaterials frequently yield only modest gains over single-function dressings and are evaluated in preclinical systems of considerable complexity^11^. Their multiple components (for example, peptides, nanoparticles, or cells) can also complicate manufacturing and the regulatory path to approval. It is therefore important to evaluate a base biomaterial separately from the benefit of functional additives, and simple, scalable, pro-regenerative dressings that also control infection are still needed.

Our approach builds on poly(polyethylene glycol citrate-co-N-isopropylacrylamide) (PPCN), a thermoresponsive, citrate-based hydrogel that can be prepared under cGMP conditions and scaled. PPCN has antioxidant properties^15,16^ and reported antibacterial properties in vitro^17^. Functionalizing PPCN with the laminin-derived peptide A5G81 promotes engagement of integrins α3β1 and α6β1 in skin, improving dermal fibroblast and keratinocyte migration and accelerating closure of full-thickness wounds in diabetic mice and swine^18,19^. In a rodent critical limb ischemia model, A5G81-PPCN also improves endothelial cell survival and limb perfusion^20^. Entrapping copper metal-organic framework nanoparticles (CMOF) in PPCN allows slow release of Cu²⁺, which limits copper toxicity and enhances vascularization and wound closure in diabetic mice^21^. These wound studies, however, were performed in non-infected wounds.

Herein, we compared four PPCN formulations (PPCN, PPCN-CMOF, A5G81-PPCN, and A5G81-PPCN-CMOF) in *S. aureus*-infected wounds in diabetic mice, using wound closure as the primary outcome. Specifically, we evaluated antibacterial activity, whether intrinsic to PPCN or arising from A5G81 or the slowly released Cu²⁺; angiogenesis, which had not previously been assessed *in vivo* across all four formulations; and granulation tissue formation, and we asked whether added functional components provided benefit beyond PPCN alone. We show that all four formulations accelerated closure of infected wounds, and that A5G81 and CMOF conferred formulation-specific differences in antibacterial activity, endothelial tube formation, vascular marker expression, and tissue architecture.

## Materials and methods

### 2.1 Materials

Citric acid, polyethylene glycol (PEG), glycerol 1,3-diglycerolate diacrylate N-isopropylacrylamide monomer (NIPAAm), 2,2-Azobisisobutyronitrile (AIBN), phosphate buffered saline (PBS), sodium hydroxide and diethyl ether were obtained from Sigma-Aldrich (St. Louis, MO). Human umbilical vein endothelial cells (HUVECs) and Human Dermal Fibroblasts (HDF) and all cell culture media were from ATCC (Manassas, Virginia). All cells, between passage 2 to 4, were cultured in their corresponding optimized culture media, in a humidified incubator equilibrated with 5% CO2 at 37°C. Tissue culture supplies were ordered from Corning Inc. (Corning, NY).

### 2.2 Preparation of A5G81-PPCN-CMOF

#### i. Synthesis of PPCN

PPCN was synthesized as previously described by our group^15^. Briefly, the poly(polyethylene glycol citrate) acrylate (PPCac) prepolymer was prepared via polycondensation by melting citric acid, polyethylene glycol (PEG), and glycerol 1,3-diglycerolate diacrylate in a 5:9:1 molar ratio at 140°C for 45 minutes under constant stirring. The resulting PPCac was then mixed with N-isopropylacrylamide (NIPAM) at a 1:1 mass ratio in 1,4-dioxane in a three-necked flask. Azobisisobutyronitrile (AIBN) was added as a radical initiator, and the reaction was carried out for 16 hours at 65 °C under a nitrogen atmosphere. The resulting product was dissolved in 1,4 dioxane, purified by precipitation in diethyl ether, and vacuum-dried to yield PPCN. The 1HNMR was performed at ambient temperature using an X500 NMR spectrometer (Bruker, Billerica, MA). Dimethyl sulfoxide (DMSO-*d6*) was used as the solvent and tetramethylsilane (TMS) as the internal reference. Attenuated Total Reflection-Fourier Transform Infrared (ATR-FTIR) spectra was obtained using the ThermoNicolet Nexus 870 spectrometer, spectra were collected with 8cm^-1^ spectral resolution and the accumulation on 64 scans. The reflectance element utilized was a ZnSe crystal, which provided ten internal reflections.

#### ii. Synthesis of A5G81-PPCN

PPCN (200 mg/mL) in 4mL of MQ water was dissolved and subsequently conjugated to N-β-maleimidopropionic acid hydrazide (BMPH, ThermoFisher, Waltham, MA)^18^, a heterobifunctional linker containing hydrazide and maleimide functional groups. 80mg of EDC and 120mg of NHS were dissolved in 1 mL MQ water, respectively and added into the PPCN solution. After a 2hrs reaction at room temperature, BMPH (24 mg) was dissolved in 1 mL of MQ water and then added to the solution to react overnight at room temperature. The PPCN-BMPH product was neutralized to pH 7.4, dialyzed using 2kDa molecular weight-cutoff (MWCO) dialysis tubing for 2 days at 4 °C, and subsequently lyophilized for 2 days. Ellman’s assay was utilized to quantify the BMPH conjugated on PPCN. PPCN and PPCN-BMPH were dissolved in PBS at 50 mg/mL. L-cysteine with concentrations ranging from 0 to 1.5 mmol was used as a standard curve. A volume of 30μL of PPCN and BMPH-PPCN was diluted in 150μL PBS, and 20μL of L-cysteine solution at 1.5 mmol was added. After adding 4μL of the Ellman’s solution, the solution was incubated at room temperature for 20min, and the absorbance was measured at 412nm using a Cytation5 plate reader. The amount of BMPH was calculated by the difference in absorbance between PPCN and BMPH-PPCN. For peptide conjugation, BMPH-PPCN (0.01 mmol) was mixed in MQ Water to react overnight at room temperature. Subsequently, the PPCN-A5G81 were dialyzed using 3.5kDa MWCO dialysis tubing for 2 days at 4 °C and lyophilized for 2 days. The amount of peptide conjugated on PPCN was quantified by TNBSA assay following the manufacturer’s protocol. PPCN and BMPH-PPCN were dissolved in 0.1 NaHCO_3_ solution at 25 mg/mL. L-glutamic acid with concentrations ranging from 0 to 1.5 mmol was used to create a standard curve. A volume of 25μL of the 0.01% (w/v) solution of TNBSA was mixed with 50μL of PPCN and BMPH-PPCN solution. After incubating at 37 °C for 2hrs, 25μL of 10% SDS solution and 12.5μL of 1M hydrochloric acid were added to the solution, and the absorbance of the resulting solution was measured at 335 nm.

#### iii. Preparation of PPCN-CMOF

Copper metal-organic framework (CMOF) synthesis followed an adapted solvothermal procedure outlined from Carson et al.^22^ Briefly, copper (II) nitrate trihydrate (1.053 g) and terephthalic acid (0.754 g) were dissolved in 87 mL of dimethylformamide (DMF) in a round-bottom flask and heated at 125°C for 24 hours at 300 rpm. The resulting blue Cu(tpa)·(dmf) crystals were precipitated with diethyl ether and repeatedly washed until the solution turned into a homogeneous turquoise. Finally, the powder was then dried under ambient conditions and stored in a desiccator. For the fabrication of PPCN-CMOF and A5G81-PPCN-CMOF (100 mg/mL), PPCN and A5G81-PPCN were loaded with 1 mM of copper-based metal-organic framework nanoparticles (CMOF), a concentration previously determined to be non-cytotoxic. The synthesis of CMOF nanoparticles was confirmed using the STADI-P X-ray diffraction (XRD) (STOE & Cie GmbH, Darmstadt, Germany), dynamic light scattering (DLS) using the Zetasizer Ultra (Malvern, UK), and scanning electron microscopy (SEM) using the Hitachi S3400N-II.

### 2.3 Rheology

The viscoelastic properties of PPCN, PPCN-CMOF, A5G81-PPCN, and A5G81-PPCN-CMOF hydrogels were evaluated using a Discovery Hybrid Rheometer DHR-3 (TA Instruments, New Castle, DE). Measurements were taken using a temperature ramp test over a temperature range of 15–45 °C with a heating rate of 2 °C/min, at a frequency of 1.5 Hz and 1.5% strain. Storage modulus (G′) and loss modulus (G″) were recorded to assess the thermoresponsive behavior. To calculate the Lower critical solution temperature (LCST), the data was plotted G’/G’’ versus Temperature and the intersection of the slope is the LCST.

### 2.4 Cytotoxicity assay

Cells were seeded in 24-well plates (5 × 10⁴ cells/well) and allowed to attach overnight. Samples of all formulations (100 µL/well) were applied using transwells (Corning, NY) and incubated overnight. Alamar Blue reagent was added according to the manufacturer’s protocol, after 4hrs 100µL of the solution was transferred to a 96-well plate and absorbance was measured at 570 nm using a Cytation5 plate reader (Agilent BioTek, Vermont, USA). A blank was included in the 24-well plate and transferred into the 96-well plate to serve as background. Cell viability was expressed as a percentage of untreated controls. For the 72-hour assay, transwells were replaced at each time point following absorbance measurement. The following equation was used to calculate viability:

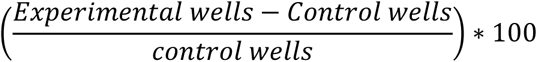

### 2.5 Antioxidant assay

The antioxidant properties of PPCN, PPCN-CMOF, A5G81-PPCN, and A5G81-PPCN-CMOF (100mg/mL) were assessed via 2,2′-azino-bis(3-ethylbenzthiazoline-6-sulfonic acid) (ABTS). A stock solution of 7 mM ABTS and 2.45 mM sodium persulfate in MQ water was prepared and left for 16 h in the dark at room temperature, after which the solution was sequentially filtered by 0.45 um filters. This working solution was then exposed to samples and incubated at 37 °C. At each timepoint (2, 8, 24, and 48 hrs), ABTS solution was sampled, and diluted with MQ water 1:1, and the absorbance was measured at 734 nm. Ascorbic acid served as the positive control and PTFE as the negative control. All measurements were performed in triplicate. The free radical scavenging activity was measured as % inhibition of free radicals by measuring the decrease in absorbance compared to control solutions

### 2.6 Antimicrobial Assay

*Staphylococcus aureus* (S. aureus) was cultured in LB broth at 37 °C and 200 rpm overnight. The bacteria were harvested, washed, and resuspended in PBS at a concentration of 1 × 10^5^ CFU/mL. A total of 100 μL of PPCN, PPCN-CMOF, A5G81-PPCN, and A5G81-PPCN-CMOF (100mg/mL) (prepared in PBS) was added to each well of a 96-well plate, followed by addition of the bacterial suspension. The plate was then incubated at 37 °C for 24 h. After incubation, 100 μL of the bacterial suspension was collected from each well and serially diluted in sterile PBS. A volume of 20 μL from each diluted sample was spread onto LB agar plates and incubated at 37 °C overnight. Digital images of the agar plates were acquired, and the colony-forming units (CFU) were quantified manually. All experiments were conducted in triplicate (n = 3). The bacterial killing ratio as calculated as follows:

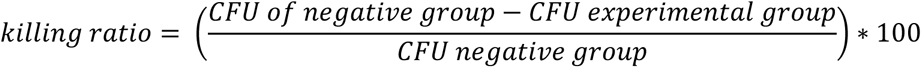

### 2.7 Tubulogenesis assay

HUVECs (2.5 × 10⁴ cells/well) were seeded in 24-well plates and allowed to adhere overnight. Cells were then treated with PPCN, A5G81-PPCN, PPCN-CMOF, and A5G81-PPCN-CMOF for 24 hours. Matrigel (diluted 1:1 with EBM) was dispensed (60 µL/well) into a 96-well plate and preincubated at 37 °C for 30 minutes. Treated cells (2 × 10⁴/well) were then plated and incubated for 4 hours. Tube formation was imaged and quantified using the Angiogenesis Analyzer plugin in ImageJ (National Institutes of Health, Bethesda, MD, USA).

### 2.8 *In vivo* studies

All animal procedures were approved by the Institutional Animal Care and Use Committee (IACUC) at Northwestern University (Protocol IS00018748). A total of 20 B6.BKS(D)-Leprdb/J (Jackson Laboratory) db/db diabetic mice were purchased for the non-infected study; however, two mice died following surgery and were excluded from further analysis, resulting in a final sample size of 18 mice. The infected study included a total of 16 mice. Full-thickness excisional wounds were created as previously described^18^. Briefly, two 6-mm circular full-thickness wounds were generated on the dorsal flanks of each mouse and stabilized with silicone splints. Wounds were treated with 40 µL of PPCN, PPCN-CMOF, A5G81-PPCN, A5G81-PPCN-CMOF, or saline, with saline-treated wounds serving as the comparator standard of care (SOC) and covered with Tegaderm. For the infected study, wounds were inoculated with *Staphylococcus aureus* (1 × 10⁵ CFU) at the time of wounding, and treatment was initiated 3 days after infection. Wound closure was monitored every 3 days. Wound area was measured from digital images using ImageJ version 1.54p (National Institutes of Health, Bethesda, MD, USA), with the inner 10 mm diameter of the splint used as the calibration reference. Percentage wound closure was calculated using the following formula:

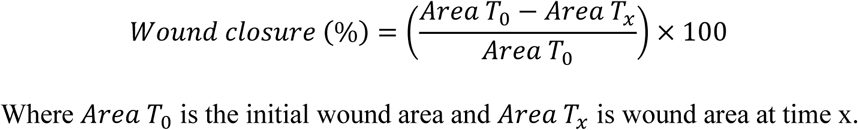

Where *Area T*_0_ is the initial wound area and *Area T_x_* is wound area at time x.

### 2.9 Histological studies

On day 27 post-surgery, all animals were euthanized, and the wounded tissue was excised and fixed in 10% formalin (Epredia, Kalamazoo, MI, USA) for 48 hours. The tissue was then washed with PBS. After fixation, the tissue was dehydrated in gradually ascending ethanol solutions from 50% to 100% (twice in 100%) and then cleared in xylene and paraffin. Subsequently, the tissue was embedded in paraffin and were sectioned 5-μm thickness onto microscope slides. Tissue sections were then stained for Masson’s trichrome staining (MTS). The tissue sections were also immunochemically stained by endothelial cell marker, CD31, and *α*-Smooth muscle actin (*α*-SMA). Briefly, the slide sections underwent deparaffinization, followed by rehydration. The rehydrated tissue sections were subjected to heat-induced epitope retrieval in pH 6 citrate buffer. Once they cooled to room temperature, the tissue sections were washed three times with PBS and subsequently blocked with a 5mg/mL bovine serum albumin solution for a duration of 30 min. Primary antibodies IgG against CD31 (1:200 clone EPR17259; catalog No. ab182981, Abcam), *α*-Smooth muscle actin(1:1000 clone 1A4; catalog No. ab7817, Abcam) were applied on tissue sections and incubated overnight at 4°C. After rinsing with PBS for 5 min three times, the fluorescently labelled secondary antibodies were applied and incubated for 30 min at room temperature. The tissue sections were then rinsed with PBST buffer for 5 min six times. Tissue was imaged using a NanoZoomer slide scanner (Hamamatsu Photonics, Shizuoka, Japan). Granulation tissue thickness was measured at five evenly spaced points across each wound using NDP.view2 software. At least 3 sections per animal were used for all the quantification.

### 2.10 Statistical methods

All data were expressed as mean ± standard deviation (SD) (n = 3 for all the in vitro studies, n = 4 for ABTS assay, and, for the in vivo studies, n = 6–8 animals per group for wound closure for a total of 36 wounds for non-infected animals and 32 wounds for infected animals, n = 6 animals per group for histology, and n = 4 per group for immunofluorescence). For the *in vitro* studies, statistical comparisons between groups were performed using one-way ANOVA followed by Tukey’s post hoc test for all pairwise comparisons; for the ABTS assay, which was analyzed separately at each time point, comparisons against the PTFE control were made using Dunnett’s test. Longitudinal wound healing was analyzed using hierarchical linear mixed-effects models to account for the randomized paired within-animal design, in which each mouse received two wounds assigned to different treatments. For histology, five measurements per wound were averaged to give one value per animal, which was the unit of analysis for all histological statistics. Treatment, time, and the treatment × time interaction were modeled as fixed effects, with random intercepts for mouse and wound nested within mouse to account for within-animal and repeated within-wound correlations. Overall healing was quantified as study-duration-normalized AUC, expressed as average wound area closed (%), and compared among groups using mixed-effects models with treatment as a fixed effect and mouse as a random effect. Comparisons of each formulation versus saline were adjusted using Dunnett’s method and comparisons among formulations were adjusted using the Holm method. Time to ≥95% wound closure was analyzed by log-rank test, with Holm-adjusted pairwise log-rank tests against saline. Data are presented as mean ± SEM for longitudinal wound closure and as box-and-whisker plots with individual data points for normalized AUC. Two-sided P < 0.05 was considered significant. In vitro and in vivo data were analyzed using GraphPad Prism 10. Statistical significance is denoted as follows: *p<0.05, **p<0.01, ***p<0.001, and ****p<0.0001.

## Results

### 3.1 A5G81-PPCN-CMOF is cell compatible, slowly releases Cu²⁺ and maintains thermoresponsive properties

PPCN and A5G81-PPCN were successfully synthesized as previously described^16^. The chemical compositions of PPCN and A5G81-PPCN were confirmed by Fourier-transform infrared spectroscopy (FTIR) and MALDI TOF (Figure S1 and S2, Supporting Information). CMOF nanoparticles were synthesized and had an average hydrodynamic diameter of 377 ± 97 nm by dynamic light scattering (DLS) (Figure 1F). Because the particles are non-spherical and aggregate (Figure 1B), DLS gives an approximate, equivalent-sphere size and only a rough estimate of polydispersity. Scanning Electron Microscopy (SEM) was therefore used to examine morphology and not to determine size. Their crystalline structure was examined by X-ray diffraction (XRD; Cu Kα, λ = 1.5406 Å) (Figure 1D), which showed principal reflections at 10.2°, 12.1°, 17.2° and 24.9° 2θ, with weaker reflections at 13.6°, 17.8°, 20.5°, 34.2° and 42.2°. These positions agree with the Cu-tpa (copper terephthalate) pattern reported by Carson et al.^22^, supporting the formation of a crystalline phase. Particle morphology was examined by SEM (Figure 1B). The Cu²⁺ concentration in A5G81-PPCN-CMOF was set at 1 mM based on cell viability measured at several copper concentrations (Figure S3, Supporting Information). Release was fastest early on, with 0.75 ± 0.19 µg/mL released by 8 hours (about half of the 72-hour total), followed by a slower phase reaching 0.96 ± 0.28 µg/mL at 24 hours, 1.20 ± 0.36 µg/mL at 48 hours and 1.46 ± 0.46 µg/mL at 72 hours (Figure 1E). The hydrogel exhibited a lower critical solution temperature (LCST) of 25 °C, which is comparable to that of PPCN alone (29 °C), indicating that copper incorporation did not significantly alter the thermoresponsive behavior (Figure 1G and 1H). The LCST was defined as the crossover point of G′ and G″. Moreover, the inclusion of copper did not reduce the viability of human dermal fibroblasts, which remained at 107 ± 16% after 72 hours (Figure 1I).

**Figure 1.**
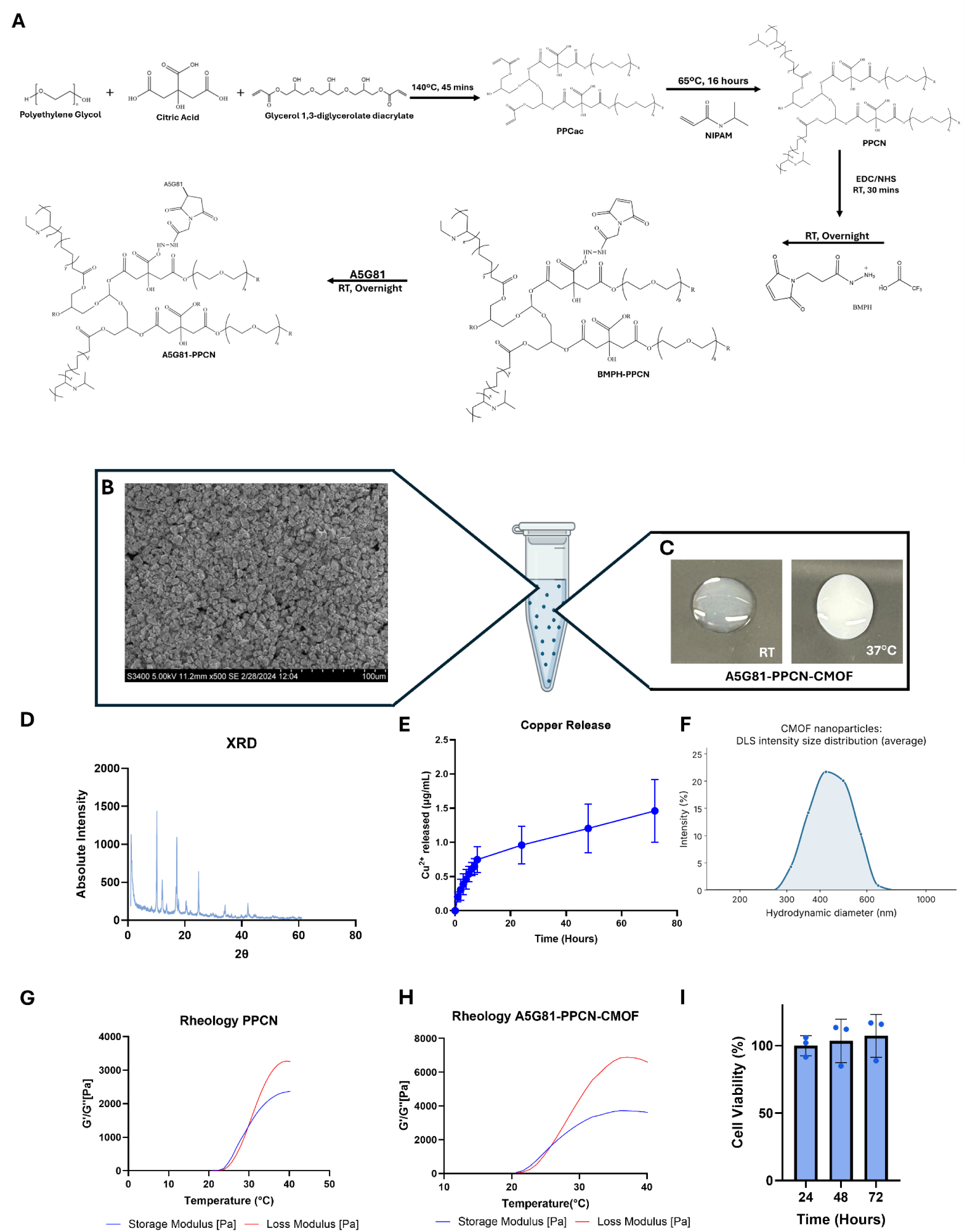
Synthesis and characterization of A5G81-PPCN-CMOF. (A) Schematic of synthesis of A5G81-PPCN. (B) SEM image of CMOF particles. (C) Digital images of A5G81-PPCN-CMOF at room temperature (left) and 37°C (right), (D) X-Ray Diffraction (XRD) of CMOF, (E) Copper release profile of A5G81-PPCN-CMOF (F) Dynamic light scattering (DLS) particle size distribution of CMOF in PBS at 25 °C. (G) Rheological characterization of PPCN showing the profiles for storage modulus G’ and loss modulus G’ (H) Rheological characterization of A5G81-PPCN-CMOF showing the profiles for storage modulus G’ and loss modulus G’’, and (I) Viability of cells exposed to A5G81-PPCN-CMOF.

### 3.2 PPCN-based hydrogels have in vitro antioxidant capacity and formulation-dependent antibacterial activity

To evaluate the antibacterial efficacy of the hydrogels, they were tested against *Staphylococcus aureus* (S. aureus), a common pathogen implicated in chronic wound infections. Penicillin-streptomycin (PS) served as the positive control, while the negative control was used as the reference for calculating the relative bacterial killing ratio. The CMOF-containing formulations exhibited the highest killing ratios, reaching 90.4 ± 6.7% for A5G81-PPCN-CMOF and 84.5 ± 2.7% for PPCN-CMOF, with no significant difference between the two formulations (p = 0.635; Figure 2B–C). In comparison, formulations without CMOF exhibited lower killing ratios of 75.6 ± 2.8% for A5G81-PPCN and 58.8 ± 9.1% for PPCN. One-way ANOVA across the four formulations showed significant differences (p = 0.001), and Tukey’s post hoc test showed that PPCN was significantly lower than PPCN-CMOF (p = 0.003), A5G81-PPCN-CMOF (p < 0.001) and A5G81-PPCN (p = 0.036). The remaining comparisons did not differ significantly: A5G81-PPCN vs PPCN-CMOF and A5G81-PPCN-CMOF vs A5G81-PPCN, whereas A5G81-PPCN-CMOF tended to be higher (p = 0.062).

**Figure 2.**
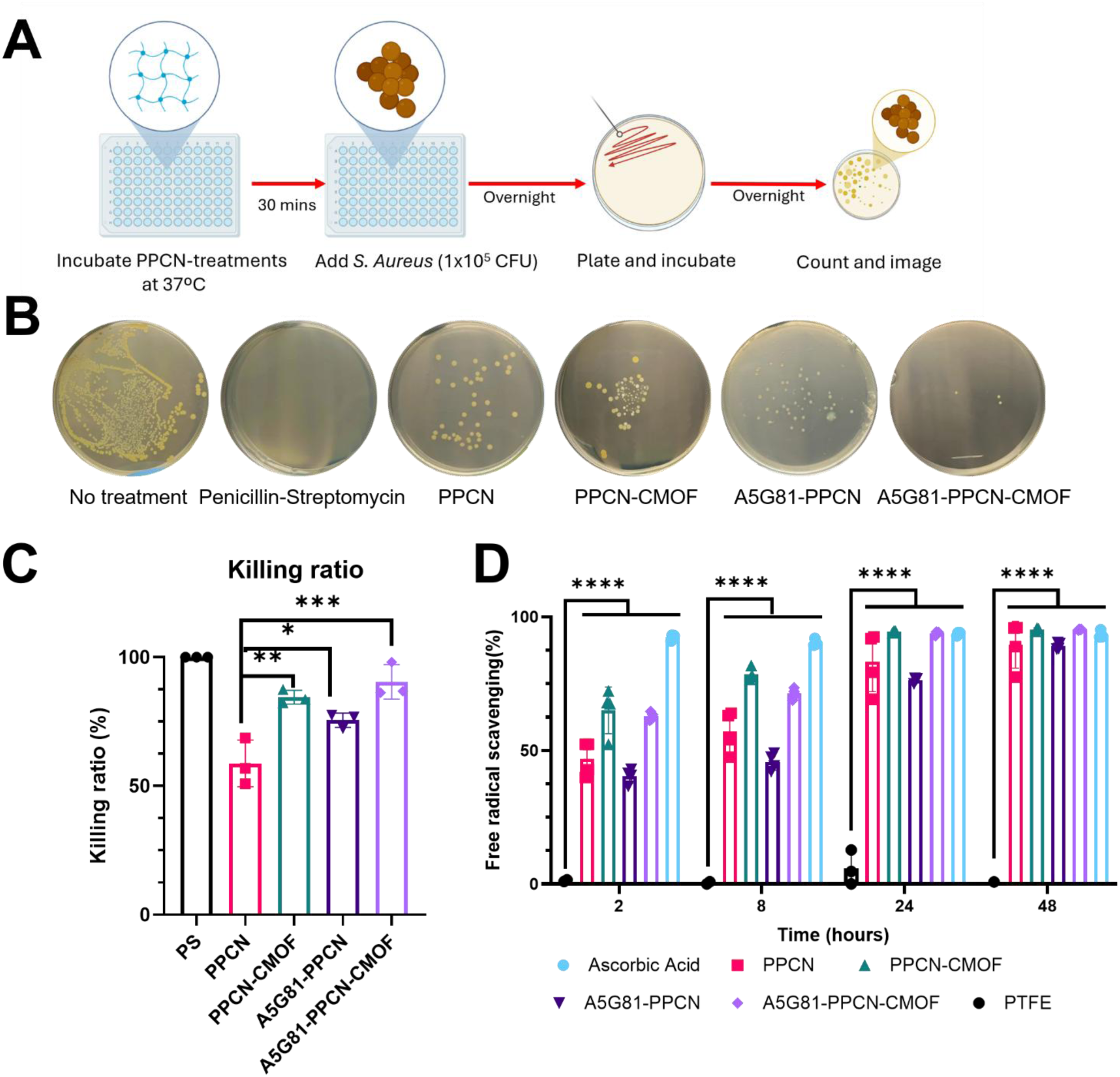
Antimicrobial and antioxidant properties of PPCN-based formulations. (A) Schematic illustration of the antimicrobial assay. (B) Representative images of bacterial culture plates following treatment. (C) Quantification of colonies imaged in B. Data are mean ± SD, n = 3; one-way ANOVA with Tukey’s test; PS excluded from testing. Penicillin–Streptomycin served as the positive control. (D) Antioxidant capacity of PPCN-based formulations, normalized to the antioxidant activity of ascorbic acid (100%). Data are presented as mean ± SD. Statistical significance is indicated as *p < 0.05, **p < 0.01, ***p < 0.001, and ****p < 0.0001.

The antioxidant capacity of the formulations was further evaluated using the ABTS free-radical scavenging assay. All four PPCN-based formulations demonstrated significantly greater radical scavenging activity than the negative control (PTFE) at every time point (p < 0.0001, Dunnett’s test), while ascorbic acid, the positive control, maintained 90–94% scavenging activity throughout the assay. At 2 hr, CMOF-containing formulations exhibited greater radical scavenging activity than their non-CMOF counterparts, with values of 65.02 ± 8.73% for PPCN-CMOF and 62.76 ± 1.58% for A5G81-PPCN-CMOF, compared with 46.83 ± 6.47% for PPCN and 40.42 ± 2.70% for A5G81-PPCN (Tukey’s test). This trend was also observed at 8 hr. By 24 hr, PPCN-CMOF (94.65 ± 0.43%), A5G81-PPCN-CMOF (93.86 ± 0.66%), and PPCN (83.20 ± 11.17%) exhibited scavenging activities that were not significantly different from ascorbic acid (93.73 ± 0.56%), whereas A5G81-PPCN remained significantly lower (76.3 ± 0.94%; p = 0.0022). At 48 hr, all formulations approached a plateau, with scavenging activities ranging from approximately 89% to 95%. At this time point, no significant differences were observed among ascorbic acid and the four PPCN-formulations. These results indicate that all four formulations have in vitro free-radical scavenging capacity; effects on oxidative tissue damage were not assessed.

### 3.3 A5G81-PPCN-CMOF has enhanced *in vitro* tubulogenesis-inducing properties

To assess the pro-angiogenic potential of A5G81-PPCN-CMOF, a tubulogenesis assay was performed using human umbilical vein endothelial cells (HUVECs) cultured on Matrigel. As shown in Figure 3, tubule-network metrics were higher than in media control for all formulations. Relative to the media control, all formulations increased mesh, junction and node formation. For mesh formation, PPCN-treated cells showed a 3.50-fold increase (p = 0.0010), whereas PPCN-CMOF, A5G81-PPCN and A5G81-PPCN-CMOF showed 6.00-fold, 6.25-fold and 7.75-fold increases, respectively (all p < 0.0001). Junction formation differed significantly among groups (p < 0.0001), with a 3.64-fold increase for PPCN (p = 0.0005), a 3.64-fold increase for PPCN-CMOF (p = 0.0005), a 3.82-fold increase for A5G81-PPCN (p = 0.0003) and a 4.27-fold increase for A5G81-PPCN-CMOF (p < 0.0001). For node formation, overall group differences were also significant (p = 0.0020), with a 2.00-fold increase for PPCN (p = 0.0081) and 1.91-fold, 2.09-fold and 2.22-fold increases for PPCN-CMOF, A5G81-PPCN and A5G81-PPCN-CMOF, respectively (p = 0.0146, p = 0.0046 and p = 0.0020). Comparisons between treatments are found in the table supplemental information **(Table 1, Supporting Information)**.

**Figure 3.**
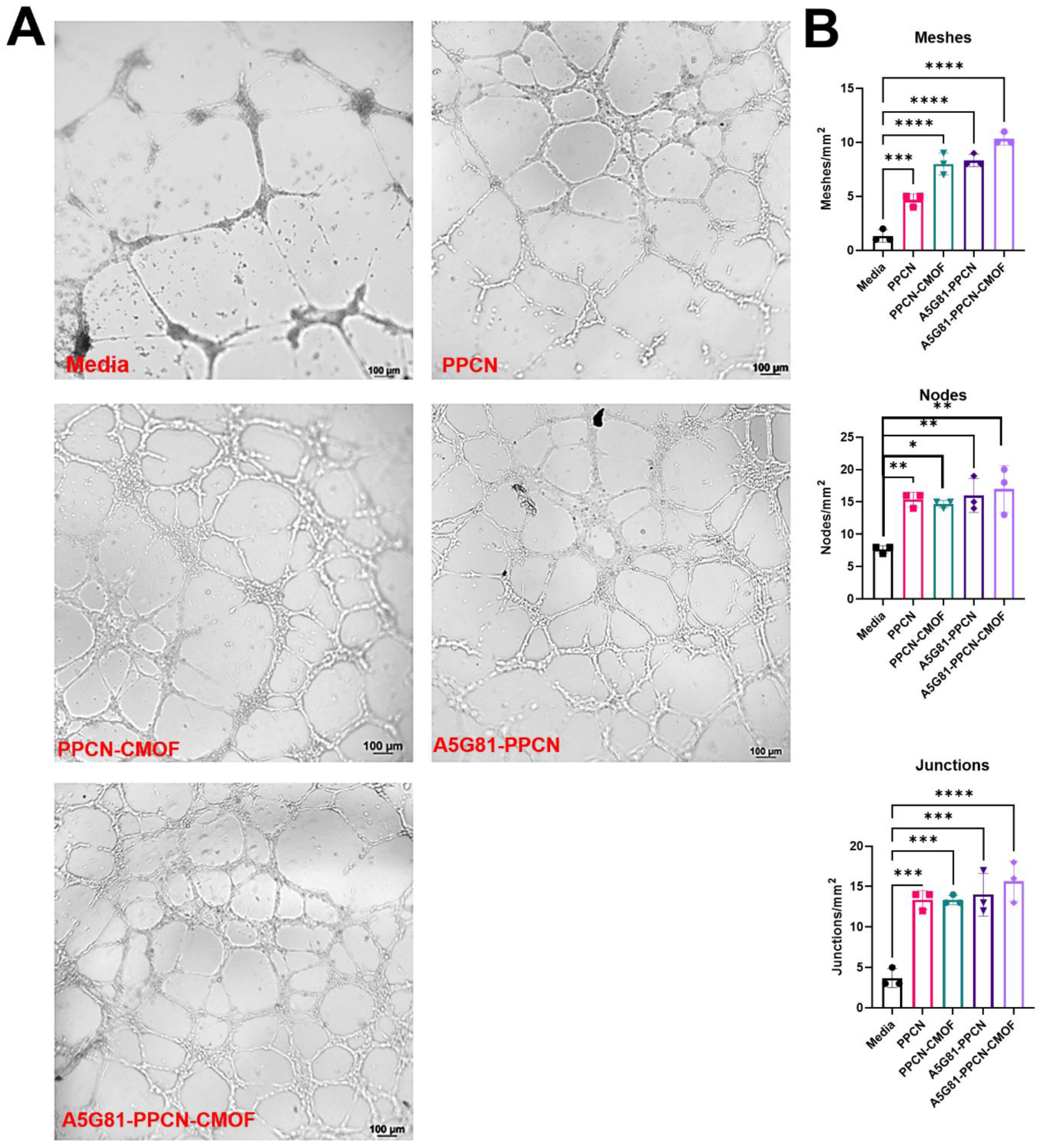
Tubulogenesis induced by PPCN-based formulations *in vitro*. (A) Representative digital images showing tubule formation by HUVECs cultured on Matrigel with media (control), PPCN, PPCN-CMOF, A5G81-PPCN, or A5G81-PPCN-CMOF. (B) Quantitative analysis of tubulogenesis, including the number of junctions, nodes, and meshes per mm², demonstrating enhanced tube network formation following treatment. Data are presented as mean ± SD (n = 3 per group); one-way ANOVA with Tukey’s test. Brackets show comparisons versus media; comparisons among formulations are reported in the text. Statistical significance is indicated as *p < 0.05, **p < 0.01, ***p < 0.001, and ****p < 0.0001.

### 3.4 In vivo wound healing efficacy of PPCN-Based formulations in a diabetic mouse model

#### 3.4.1 Effect of PPCN-based formulations on closure of non-infected full-thickness excisional wounds in diabetic mice

To evaluate the regenerative potential of our PPCN hydrogel formulations, we evaluated their capacity to heal wounds in a splinted, full-thickness excisional wound model in diabetic mice. Each animal received two dorsal wounds treated with either saline, PPCN, PPCN-CMOF, A5G81-PPCN, or A5G81-PPCN-CMOF.

In non-infected wounds, all groups approached complete closure by day 27 (Figure 4A). Cumulative healing, quantified as normalized AUC, was significantly higher than saline only for PPCN-CMOF (p = 0.046); PPCN (p = 0.60), A5G81-PPCN (p = 0.60) and A5G81-PPCN-CMOF (p = 0.33) did not differ significantly from saline (Figure 4B; n = 7, 6, 8, 7 and 8 for saline, PPCN, PPCN-CMOF, A5G81-PPCN and A5G81-PPCN-CMOF, respectively. No pairwise difference among treatments was significant after Dunnett’s method across all pairwise comparisons (all adjusted p ≥ 0.41).

**Figure 4.**
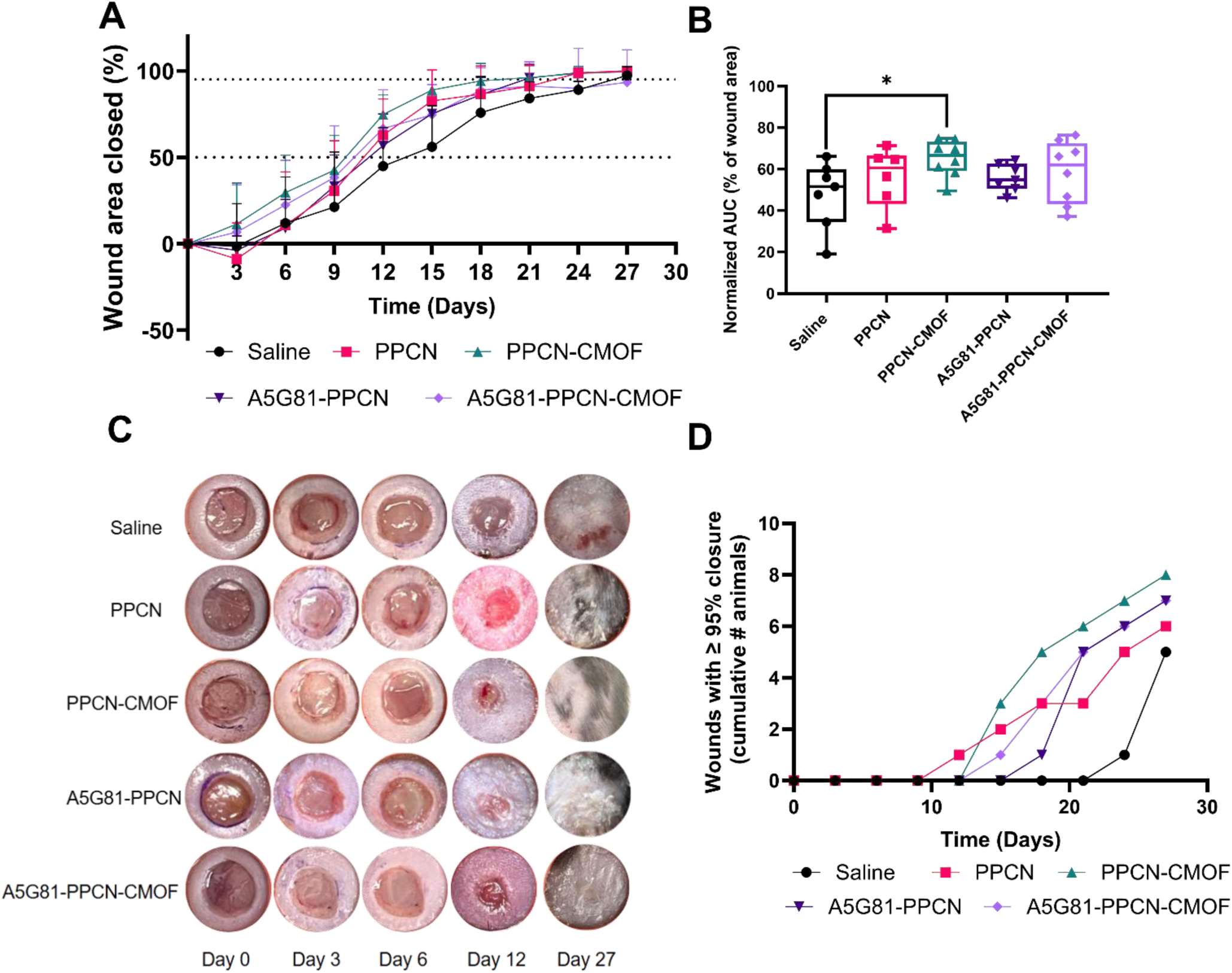
Efficacy of PPCN formulations in non-infected diabetic mice wounds. (A) Longitudinal wound closure kinetics in non-infected diabetic mouse wounds (n = 7, 6, 8, 7, and 8 wounds for saline, PPCN, PPCN-CMOF, A5G81-PPCN and A5G81-PPCN-CMOF, respectively). (B) Cumulative healing quantified as study-duration-normalized AUC (days 0–27) and expressed as average wound area closed (%). *P* values are Holm-adjusted comparisons versus saline (four comparisons): \**P* < 0.05. No pairwise comparison between treatment groups was significant after Holm adjustment. (C) Digital images of the wounds. (D) Time to ≥ 95% wound closure, all treatments reached ≥95% closure significantly earlier than saline (adjusted p = 0.009–0.045). Numbers in Y-axis indicate wounds reaching ≥95% closure / total wounds per group. Groups were compared by log-rank test (p = 0.023), with pairwise log-rank tests against saline adjusted by Holm’s method.

To examine the impact of our treatment on the quality of regenerated tissue, we performed histological analyses of collagen content and skin architecture using Masson’s trichrome staining (Figure 5). Epidermal thickness was 54.47 ± 1.271 μm in the saline control group. Treatment with PPCN alone resulted in a comparable epidermal thickness of 60.98 ± 6.335 μm, while the PPCN-CMOF and A5G81-PPCN groups exhibited mean epidermal thicknesses of 64.51 ± 9.116 μm and 73.39 ± 7.394 μm, respectively. A5G81-PPCN-CMOF treatment produced the greatest epidermal thickness, reaching 182.4 ± 43.39 μm, a significant increase compared with the saline, PPCN, PPCN-CMOF and A5G81-PPCN groups (p < 0.0001) (Figure 5B). Granulation tissue thickness was 420.16 ± 27.47 μm, 337.25 ± 8.60 μm, 562.14 ± 49.98 μm, and 563.75 ± 47.55 μm for PPCN, PPCN-CMOF, A5G81-PPCN, and A5G81-PPCN-CMOF, respectively. Saline-treated wounds had a mean granulation tissue thickness of 252.02 ± 49.38 μm, which was significantly lower than that of PPCN, A5G81-PPCN and A5G81-PPCN-CMOF (p < 0.0001) and PPCN-CMOF (p < 0.001) (Figure 5C). These results indicate greater granulation tissue thickness with all PPCN-based formulations, and greater epidermal thickness with A5G81-PPCN-CMOF, at day 27 in non-infected diabetic wounds.

**Figure 5.**
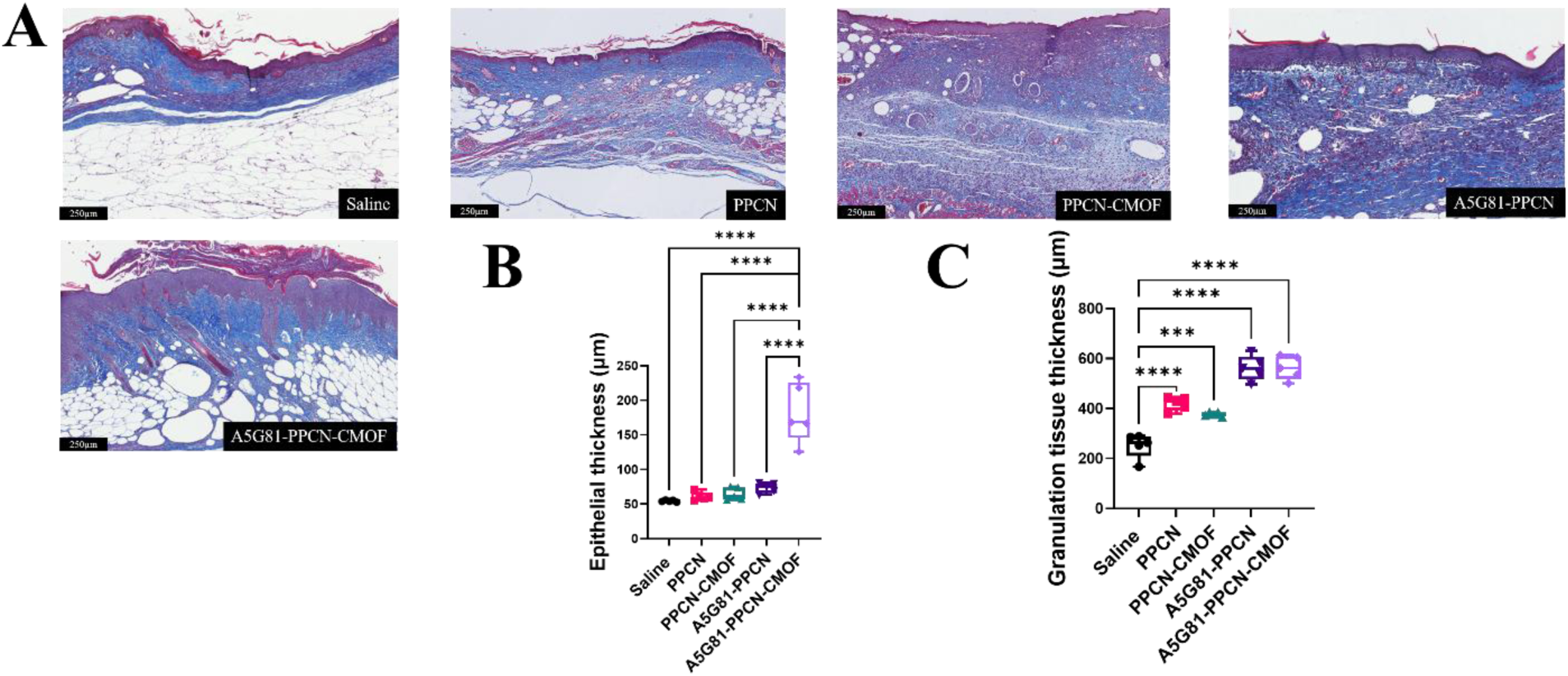
Histological assessment of wound healing following treatment. (A) Representative histological images of wound tissue collected on day 27 post-wounding. (B) Quantification of epidermal thickness. (C) Quantification of granulation tissue thickness. Data are presented as mean ± SD. Statistical significance is indicated as *p < 0.05, ****p < 0.0001. Each data point represents one animal (mean of five measurements per animal); (n = 6 animals per group).

#### 3.4.2 Infected full-thickness excisional wounds treated with PPCN-based formulations exhibit accelerated healing in diabetic mice

To evaluate the therapeutic efficacy of PPCN-formulations on infected conditions, we used a full-thickness excisional wound model in diabetic mice (n = 6–7 per group). Each animal received two dorsal wounds, and both were inoculated with *S. aureus* at the time of injury. Treatment with saline, PPCN, PPCN-CMOF, A5G81-PPCN, or A5G81-PPCN-CMOF was initiated on day 3 post-bacterial inoculation. In our model, biofilm formation was observed between days 3 and 6 post-injury in infected wounds. Notably, by day 9, biofilm in CMOF-treated infected wounds produced a red pigment, an observation that was not quantified (Figure 6C). By day 12, PPCN, PPCN-CMOF and A5G81-PPCN-treated wounds had each about 45% closure, compared with 21% for A5G81-PPCN-CMOF and 16% for saline-treated wounds, which had exhibited a −18.4% in wound closure on day 3, indicating wound expansion likely due to infection-related complications. Cumulative healing, quantified as normalized AUC, differed among treatments. Relative to saline (0.23 ± 0.05, estimated marginal mean ± SE), normalized AUC was significantly greater for PPCN (0.49 ± 0.05, p = 0.002), A5G81-PPCN (0.44 ± 0.05, p = 0.008), PPCN-CMOF (0.43 ± 0.05, p = 0.013) and A5G81-PPCN-CMOF (0.42 ± 0.05, p = 0.021; Dunnett-adjusted; Figure 6B). No pairwise difference among treatments was significant after Holm adjustment across all pairwise comparisons (all adjusted p ≥ 0.85).

**Figure 6.**
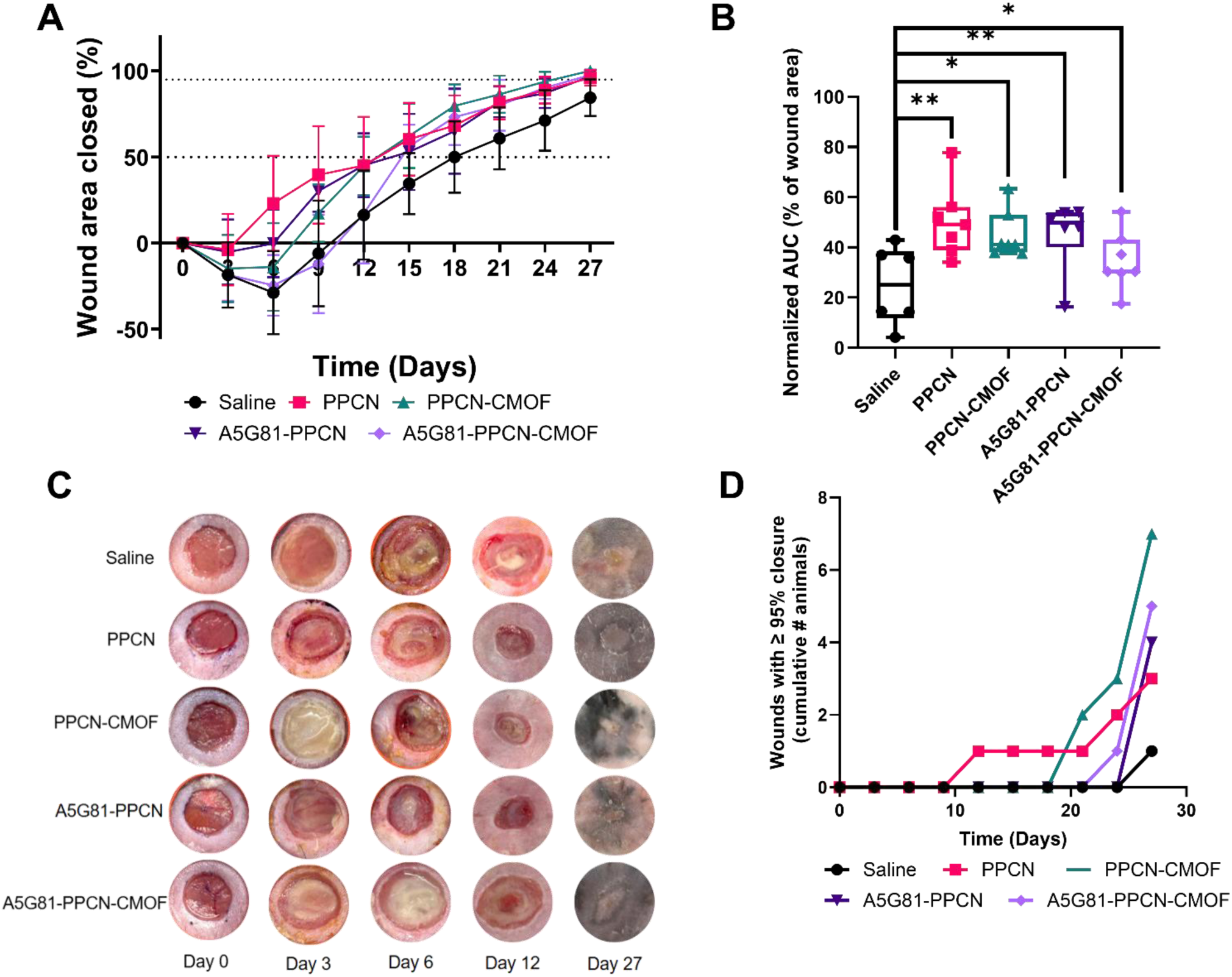
Efficacy of PPCN formulations in infected diabetic mice wounds. (A) Longitudinal wound closure kinetics in infected diabetic mouse wounds (n = 6, 7, 7, 6, and 6 wounds for saline, PPCN, PPCN-CMOF, A5G81-PPCN and A5G81-PPCN-CMOF, respectively). (B) Cumulative healing quantified as study-duration-normalized AUC (days 0–27) and expressed as average wound area closed (%). *P* values are Holm-adjusted comparisons versus saline (four comparisons): ns, not significant; \**P* < 0.05, \*\**P* < 0.01, \*\*\**P* < 0.001. Among treatment groups, only PPCN-CMOF versus PPCN differed significantly (*P* = 0.028, Holm-adjusted across all pairwise comparisons) (C) Digital images of the wounds. (D) Time to ≥ 95% wound closure. Only PPCN-CMOF differed from saline (adjusted p = 0.012). Numbers in the Y-axis indicate wounds reaching ≥95% closure / total wounds per group. Groups were compared by log-rank test (p = 0.023), with pairwise log-rank tests against saline adjusted by Dunnett’s method.

These findings were supported by histological analysis in which the mean epidermal thickness was 61.68 ± 6.179µm, 50.05 ± 6.748 μm, 77.05 ± 9.720 μm, and 54.91 ±10.25 µm for PPCN, PPCN-CMOF, A5G81-PPCN and A5G81-PPCN-CMOF groups, respectively (Figure 7). Saline-treated wounds had an epidermal thickness of 46.29 ± 3.335 µm, which was significantly lower than that of PPCN (p < 0.05) and A5G81-PPCN (p < 0.0001) (Figure 7B). Granulation tissue thickness was 510.9 ± 81.80 µm, 568.1 ± 54.80 μm, 553.8 ± 55.04 μm, and 544.33 ± 61.04 µm for PPCN, PPCN-CMOF, A5G81-PPCN and A5G81-PPCN-CMOF, respectively, compared with a mean granulation tissue thickness of 440.93 ± 43.57 µm in saline-treated wounds. The increase was significant for PPCN-CMOF, A5G81-PPCN and A5G81-PPCN-CMOF (p < 0.05) but not for PPCN (Figure 7C).

**Figure 7.**
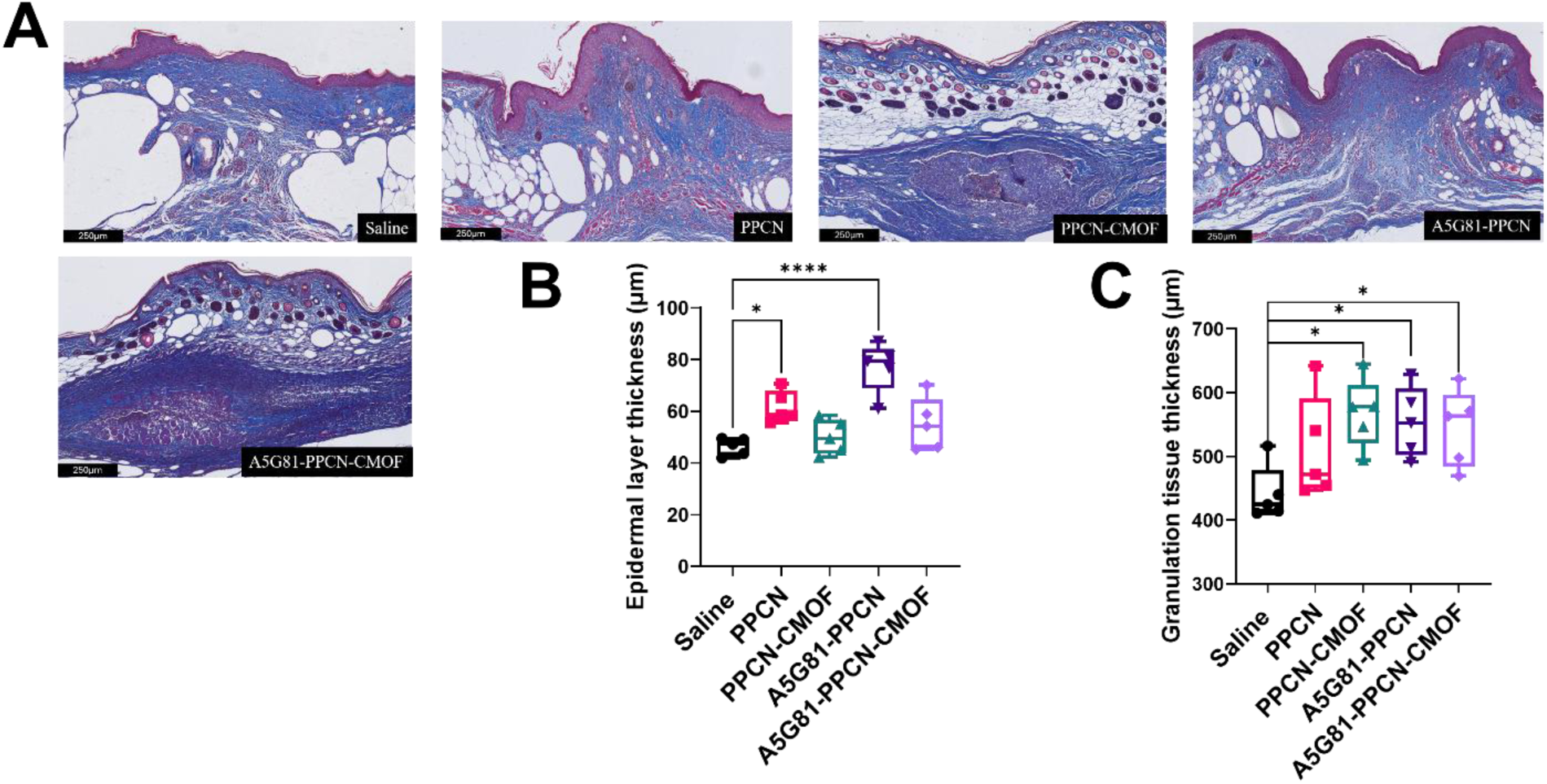
Histological assessment of infected wound healing following treatment. (A) Representative histological images of wound tissue collected on day 27 post-wounding. (B) Quantification of epidermal thickness. (C) Quantification of granulation tissue thickness. Data are presented as mean ± SD. Statistical significance is indicated as *p < 0.05, **p < 0.01, ***p < 0.001, and ****p < 0.0001. Each data point represents one animal (mean of five measurements per animal); (n = 6 animals per group).

### 3.5 Vascular and myofibroblast marker expression in non-infected and infected wounds

Immunofluorescence probing for CD31, an endothelial marker, was performed on healed tissue from non-infected mice to evaluate vascularization. Compared to the saline-treated group, PPCN groups had a 1.32-fold increase in CD31 signal, when A5G81 was tethered to PPCN there was a 1.21-fold increase, when compared to saline and a 0.92-fold change relative to PPCN alone. Moreover, PPCN-CMOF exhibited a 3.73-fold increase relative to saline and a 2.83-fold increase relative to PPCN (neither significant). The greatest CD31 signal was observed in the A5G81-PPCN-CMOF group, which exhibited a 6.82-fold increase when compared to saline (p < 0.001), a 5.18-fold increase compared to PPCN (p = 0.0019), and a 5.64-fold increase compared to A5G81-PPCN (p = 0.0016). The 1.83-fold difference relative to PPCN-CMOF was not significant (p = 0.10), so a synergistic effect of A5G81 and CMOF on endothelial-associated signaling cannot be concluded. α-SMA fluorescence staining was also evaluated, in this case, PPCN had a 0.76-fold change (a decrease) compared to saline, whereas PPCN-CMOF, A5G81-PPCN and A5G81-PPCN-CMOF showed a 2.07-fold, 2.69-fold and 1.72-fold increase, respectively (nonsignificant). Relative to PPCN, PPCN-CMOF increased 2.72-fold, A5G81-PPCN increased 3.53-fold (p = 0.030 after log transformation; p = 0.33 on untransformed data), the only α-SMA difference below p = 0.05, and A5G81-PPCN-CMOF increased 2.26-fold. Comparing PPCN-CMOF and A5G81-PPCN we observed a 0.77-fold change. Incorporating copper into A5G81-PPCN resulted in a 0.64-fold change. A5G81-PPCN-CMOF showed a 1.72-fold increase relative to saline (Figure 8). When taking the ratio of CD31: SMA, to evaluate the balance between endothelial cell signaling and myofibroblast signaling we obtained that saline was 0.309, PPCN was 0.536, A5G81-PPCN was 0.139, PPCN-CMOF was 0.557, and A5G81-PPCN-CMOF was 1.227, these descriptive ratios (not statistically tested) are consistent with a shift toward a relatively greater endothelial-associated signaling in the A5G81-PPCN-CMOF group.

**Figure 8.**
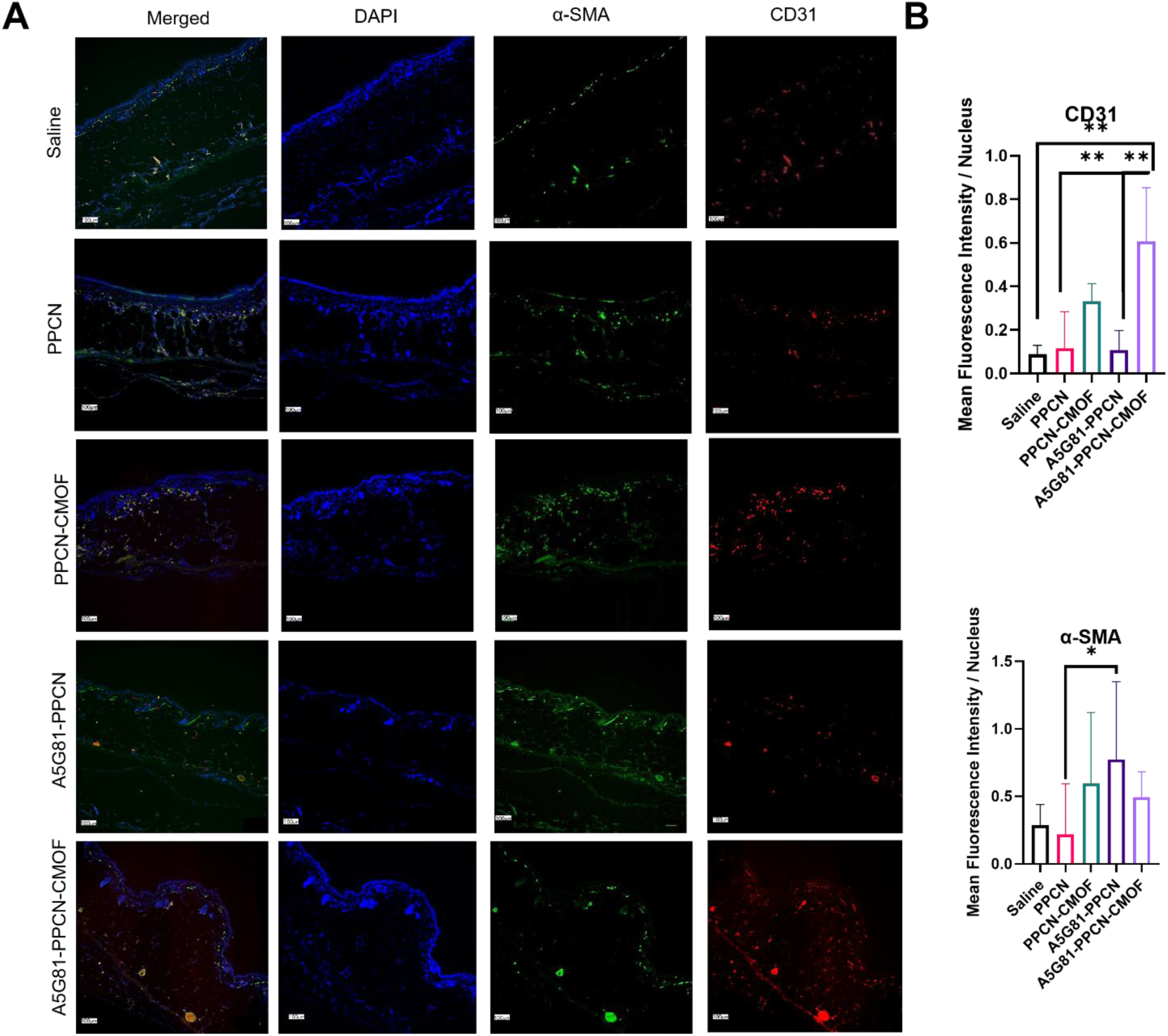
Immunofluorescence analysis of vascular markers in non-infected wound tissues treated with saline or PPCN-based formulations. (A) Representative immunofluorescence images showing merged nuclei (DAPI, blue), α-SMA (green), and CD31 (red) channels. (B) Quantification of mean fluorescence intensity normalized to nuclei count. Data are presented as mean ± SD (n ≤ 4). Unless otherwise indicated, comparisons between treatment groups were not statistically significant. Statistical significance is indicated as *p < 0.05, **p < 0.01, and ***p < 0.001.

In the infected wound model, CD31 immunofluorescence showed a 1.36-fold increase in the PPCN-treated group relative to saline, whereas PPCN-CMOF, A5G81-PPCN and A5G81-PPCN-CMOF exhibited 11.39-fold, 9.80-fold and 4.07-fold increases, respectively; only the PPCN-CMOF increase was statistically significant (p = 0.038). Relative to PPCN, PPCN-CMOF demonstrated an 8.36-fold increase in CD31 signal, A5G81-PPCN a 7.19-fold increase, and A5G81-PPCN-CMOF a 2.98-fold increase. Interestingly, incorporation of CMOF into A5G81-PPCN resulted in a 0.42-fold change relative to A5G81-PPCN, suggesting a trend toward lower CD31 signal in the copper-containing formulation under infected conditions (not significant). PPCN-CMOF and A5G81-PPCN showed comparable CD31 signal (1.16-fold), and A5G81-PPCN-CMOF showed a 0.36-fold change relative to PPCN-CMOF. None of the CD31 comparisons among formulations was statistically significant. Analysis of α-SMA staining revealed increases of 2.17-fold, 1.51-fold, 3.15-fold, and 4.35-fold for PPCN, PPCN-CMOF, A5G81-PPCN, and A5G81-PPCN-CMOF, respectively, relative to saline; only the increase with A5G81-PPCN-CMOF (p = 0.0040) was significant. Compared with PPCN, A5G81-PPCN and A5G81-PPCN-CMOF exhibited 1.45-fold and 2.00-fold higher α-SMA fluorescence, respectively (both not significant), whereas PPCN-CMOF showed a 0.70-fold change. PPCN-CMOF showed a 0.48-fold change relative to A5G81-PPCN. Addition of CMOF to A5G81-PPCN resulted in a 1.38-fold increase in α-SMA signal (not significant), and A5G81-PPCN-CMOF was 2.88-fold higher than PPCN-CMOF (p = 0.028) (Figure 9). We assessed the ratio of CD31:α-SMA for saline, PPCN, PPCN-CMOF, A5G81-PPCN, A5G81-PPCN-CMOF, which were 0.145, 0.091, 1.091, 0.450, 0.135, respectively.

**Figure 9.**
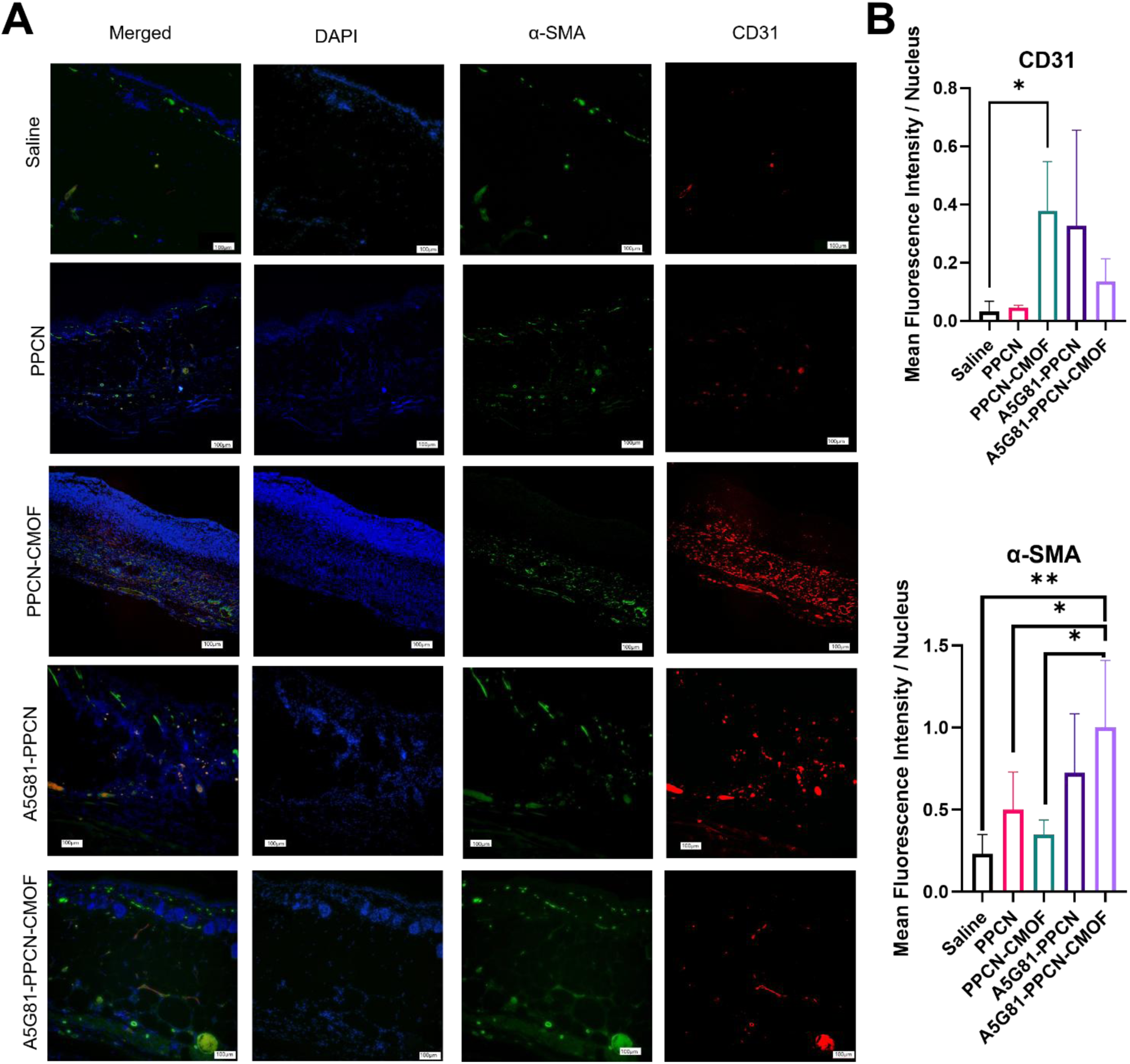
Immunofluorescence analysis of vascular markers in infected wound tissues treated with saline or PPCN-based formulations. (A) Representative immunofluorescence images showing merged nuclei (DAPI, blue), α-SMA (green), and CD31 (red) channels. (B) Quantification of mean fluorescence intensity normalized to nuclei count. Data are presented as mean ± SD (n ≤ 4). Unless otherwise indicated, comparisons between treatment groups were not statistically significant. Statistical significance is indicated as *p < 0.05 and **p < 0.01.

## Discussion

In this study, we compared four PPCN formulations in diabetic mice with non-infected and *S. aureus*-infected full-thickness wounds. The principal finding is that all four formulations significantly improved cumulative closure of infected wounds relative to saline (p = 0.002–0.021), whereas in non-infected wounds only PPCN-CMOF did (p = 0.046).

The in vitro antibacterial activity was formulation-dependent (**Figure 2**). Killing ratios relative to the untreated control were highest for the CMOF-containing formulations (90.4% for A5G81-PPCN-CMOF and 84.5% for PPCN-CMOF), intermediate for A5G81-PPCN (75.6%), and lowest for PPCN (58.8%). PPCN was significantly lower than PPCN-CMOF (p = 0.003), A5G81-PPCN-CMOF (p < 0.001), and A5G81-PPCN (p = 0.036), whereas the other pairwise comparisons were not significant (**Figure 2C**). Copper has well-described antibacterial mechanisms, including membrane damage, ROS generation, and oxidation of lipids, proteins, and DNA^15,23,24^, which is consistent with the greater killing by the CMOF-containing formulations, although the CMOF content (1 mM Cu²⁺) was selected for cytocompatibility with fibroblasts rather than for antibacterial potency. Laminin and integrin-derived peptide fragments have been implicated in host defense^25^, but a direct antibacterial role for A5G81 has not been reported. Because A5G81-PPCN-CMOF did not differ significantly from A5G81-PPCN or PPCN-CMOF, we cannot conclude that the combination is superior or synergistic. PPCN formulations also scavenged ABTS radicals in vitro, significantly more than the PTFE control at every time point (p < 0.0001), with CMOF-containing formulations scavenging more at early time points (65.02% and 62.76% for PPCN-CMOF and A5G81-PPCN-CMOF vs 46.83% and 40.42% for PPCN and A5G81-PPCN at 2 hr) and all formulations plateauing at about 89–95% by 48 hr, consistent with prior reports^17,26^, but we did not measure oxidative damage in tissue, so the contribution of antioxidant activity to healing remains undemonstrated (**Figure 2D**).

A5G81 and CMOF together influenced endothelial behavior in vitro (**Figure 3**). In HUVEC tubulogenesis assays, all formulations increased mesh, junction, and node formation relative to media up to 7.75-, 4.27-, and 2.22-fold, respectively for A5G81-PPCN-CMOF. Relative to PPCN, A5G81 increased mesh formation (1.79-fold, p = 0.0005) but not junction (1.05-fold, p = 0.99) nor node formation (1.04-fold). Further, the addition of CMOF (A5G81-PPCN-CMOF vs A5G81-PPCN) increased mesh formation (1.24-fold, p = 0.0316) but not junction (1.12-fold, p = 0.79) or node formation (1.06-fold, p = 0.97). This is consistent with reports that laminin-derived peptides enhance angiogenesis and wound repair^25,26^ and that copper promotes endothelial responses^22^. PPCN-CMOF vs PPCN increases meshes (1.71-fold, p = 0.0010) but not junctions or nodes.

In non-infected wounds, all groups, including saline, approached closure by day 27 (mean 93–100%), so differences lay mainly in healing kinetics (**Figure 4**). At day 12, closure was 57–75% for the PPCN formulations compared with 45% for saline, and all four formulations reached ≥95% closure earlier than saline (adjusted p = 0.009–0.045), although only PPCN-CMOF significantly improved cumulative closure (**Figure 4D**). The effect of PPCN-CMOF is consistent with our earlier report of CMOF-containing PPCN in diabetic mice^21^.

Whereas healing was markedly impaired in infected wounds although early closure differed among formulations (**Figure 6A**). Saline-treated wounds expanded during the first days (−18%, −29% and −6%, on days 3, 6, and 9) and reached only 16% closure at day 12 and 84% by day 27. This slowed healing is consistent with reports that elevated bioburden correlates with impaired healing^27^ and that bacteria increase inflammation, necrosis, and moisture in the wound bed, effects likely compounded by diabetic immune dysfunction^28,29^. In contrast, PPCN showed the fastest early closure (−4%, 23%, and 40% at days 3, 6, and 9) and was significantly higher than saline at days 6 and 9 (p = 0.005 and p = 0.011). PPCN was followed by A5G81-PPCN (−5%, 0%, and 30%) and PPCN-CMOF (−15%, −14%, and 17%), whereas A5G81-PPCN-CMOF remained below its day-0 area through day 9 (−17%, −23%, and −8%). On day 9, PPCN closure was significantly higher than A5G81-PPCN-CMOF (Holm-adjusted p = 0.037); By day 12, closure was about 45% for PPCN, PPCN-CMOF and A5G81-PPCN, compared with 21% for A5G81-PPCN-CMOF and 16% for saline. PPCN-CMOF showed the highest closure from day 18 onward (80%, 86%, 94%, and 100% on days 18, 21, 24, and 27), although it did not differ significantly from the other treatments in those days. Its closure was significantly higher than saline on day 15 (p = 0.027) and on days 21 and 24 (p = 0.018 and p = 0.003), but not on day 18 (p = 0.065). By day 27, mean closure in all four treatment groups was significantly higher than in saline with 96%, 100%, 97%, and 97%, for PPCN, PPCN-CMOF, A5G81-PPCN, and A5G81-PPCN-CMOF, respectively, compared with 84% for saline; (p = 0.003 for PPCN, A5G81-PPCN and A5G81-PPCN-CMOF, and p < 0.001 for PPCN-CMOF). Because this benefit was shared across formulations, it may arise at least in part from properties of the PPCN hydrogel itself, such as conformal coverage of the wound bed after thermoresponsive gelation. Our design did not include a non-PPCN hydrogel control, so we cannot separate PPCN chemistry from general hydrogel effects. Moreover, we did not quantify bacterial burden in vivo therefore we cannot attribute that the closure benefit results from bacterial killing, reduced biofilm, or infection clearance. It may instead reflect an improved host response or tissue regeneration in the presence of infection.

Tissue architecture followed a similar formulation-dependent pattern. In non-infected wounds, A5G81-PPCN-CMOF produced the thickest epidermis (p<0.0001 vs all other groups), and granulation tissue was thicker in all PPCN formulations than in saline (**Figure 5**). In infected wounds, granulation tissue was thicker with PPCN-CMOF, A5G81-PPCN, and A5G81-PPCN-CMOF, whereas epidermal thickness increased with PPCN and A5G81-PPCN but not with the CMOF formulations (**Figure 7**). These patterns may reflect formulation-and infection-dependent effects on re-epithelialization and granulation tissue formation, which require further study. Greater thickness should not be equated with better tissue quality and may partly reflect the healing stage. Measures of epithelial barrier integrity, collagen organization, and functional restoration would be needed to support improved regeneration.

In vivo the non-infected wounds treated with A5G81-PPCN-CMOF produced the highest CD31 signal, and in infected wounds PPCN-CMOF did, but the pattern depended on infection status and most comparisons were not significant (**Figures 8 and 9**). Laminin-derived peptides have been reported to promote endothelial responses, Demidova-Rice et al.^30^ demonstrated enhanced blood vessel formation in impaired cutaneous wound-healing models; Ishihara et al. found that laminin-derived peptides improved wound repair and increased angiogenesis, possibly by facilitating the release and activity of proangiogenic growth factors^31^. Our in vitro mesh-formation results are consistent with such a role for A5G81, but our in vivo data do not show that A5G81 drove the CD31 differences, and CD31 staining does not establish increased functional vessel density or perfusion.

On the other hand, α-SMA in infected wounds was highest with A5G81-PPCN-CMOF, which was significantly higher than saline and PPCN-CMOF. CD31 measures endothelial-associated staining, α-SMA labels perivascular cells as well as myofibroblasts, and the CD31 to α-SMA ratio cannot by itself indicate vascular maturation. Most comparisons were not significant at n ≤ 4. The vascular response is therefore not a simple additive effect of A5G81 and CMOF, and infection appears to change it, so we avoid concluding that the components act synergistically. Co-localization analyses and perfusion measurements would strengthen mechanistic interpretation.

An important implication is that the incremental value of functionalization should be judged against the added complexity. A5G81 and CMOF were each associated with significantly greater in vitro killing than PPCN alone, and adding CMOF produced endpoint-specific changes in endothelial network formation and tissue staining. Yet neither clearly translated into faster infected-wound closure than PPCN alone in this model; cumulative closure did not differ significantly among formulations. Given the manufacturing, safety, and regulatory burden of additional components, PPCN itself, which can be prepared under cGMP conditions and scaled, remains an attractive base platform, with peptides or nanoparticles added only where a benefit is demonstrated. Future work should quantify in vivo bacterial burden and biofilm, dissect the antibacterial contribution of A5G81, and evaluate the most promising formulations in a larger animal model, where A5G81-PPCN has already shown benefit in non-infected wounds^17^.

## Conclusion

In summary, PPCN-based hydrogels significantly improved cumulative closure of non-infected and *S. aureus*-infected wounds in diabetic mice, with benefits observed across all four formulations. A5G81 and CMOF functionalization produced distinct effects on antibacterial activity in vitro, endothelial tubulogenesis, vascular-associated markers, and tissue architecture, but these modifications did not demonstrate consistent additional benefit for wound closure relative to unmodified PPCN. These findings support the potential of PPCN as a platform for infected diabetic wounds and highlight the importance of balancing intrinsic bioactivity, functionalization-dependent benefits, and formulation complexity when developing advanced dressings. Quantifying bacterial burden in vivo and testing the most promising formulations in larger animal models are the next steps.

## Acknowledgements

The authors would like to thank Chongwen Duan and Ruyue Luo for their help during animal studies. As well as Reyhan Nur Civ for her help. Figures created with BioRender.com and Canva.com. This research is supported by the CARE Center, the NIH grant T32-EB031527, and the National Institute of Diabetes and Digestive and Kidney Diseases (5R01DK131302-02). This work made use of the Keck-II facility (RRID: SCR_026360) of Northwestern University’s NUANCE Center, which has received support from the IIN and Northwestern’s MRSEC program (NSF DMR-2308691). This work made use of the IMSERC (RRID:SCR_017874) Crystallography facility at Northwestern University, which has received support from the Soft and Hybrid Nanotechnology Experimental (SHyNE) Resource (NSF ECCS-2025633), and Northwestern University. This work made use of the IMSERC (RRID:SCR_017874) NMR facility at Northwestern University, which has received support from the Soft and Hybrid Nanotechnology Experimental (SHyNE) Resource (NSF ECCS-2025633) and Northwestern University. Elemental analysis was performed at the Northwestern University Quantitative Bulk-Elemental Information Core (RRID:SCR_017773). Northwestern University Pathology Core Facility and a Cancer Center Support Grant (NCI CA060553) for providing their histological services. This work made use of the EPIC facility (RRID: SCR_026361) of Northwestern University’s NUANCE Center, which has received support from the IIN and Northwestern’s MRSEC program (NSF DMR-2308691).

## Supplemental Information

**Figure S1.**
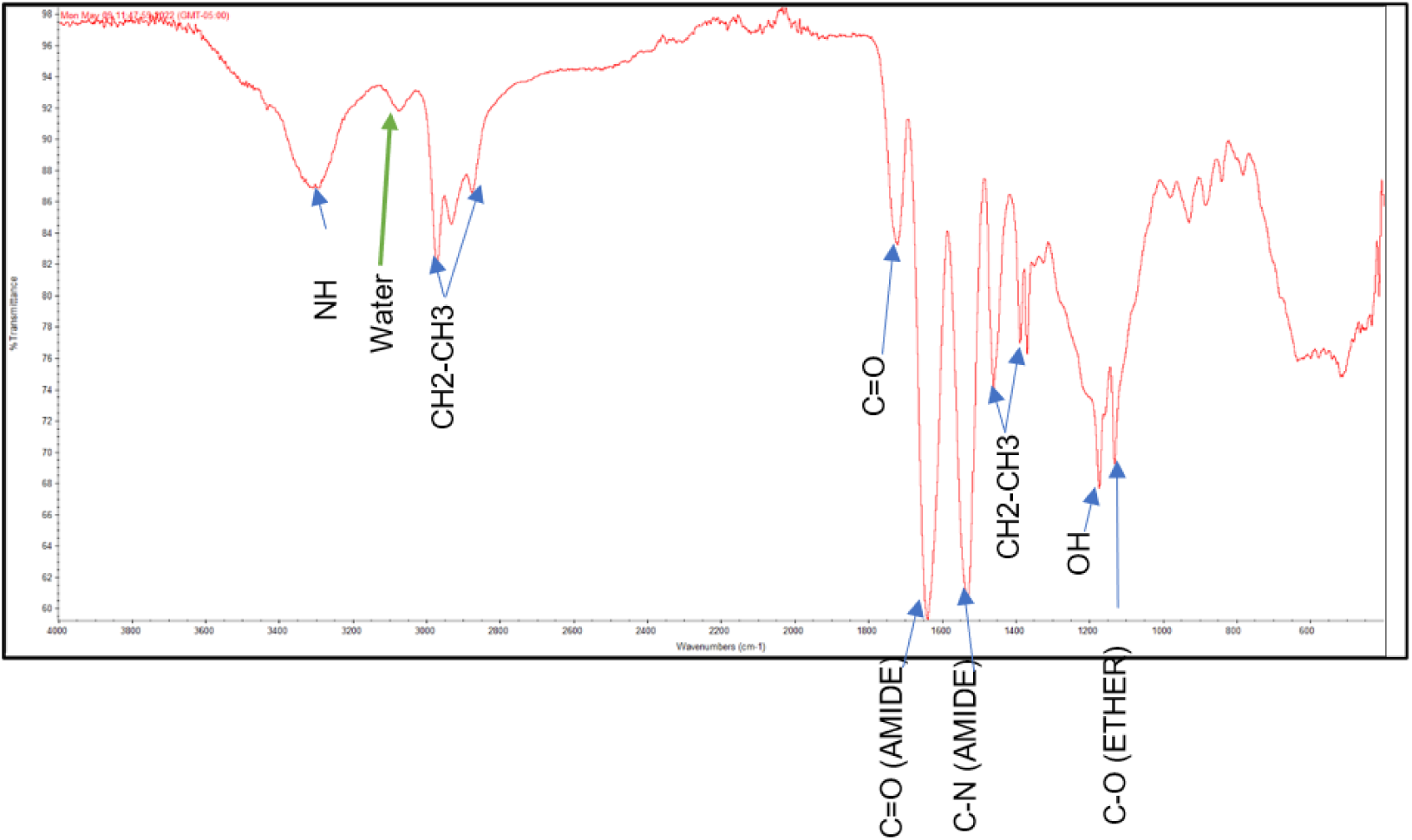
FT-IR Identification of different functional groups within PPCN. FTIR spectra confirm the formation of poly(polyethylene glycol citrate-co-N-isopropylacrylamide) (PPCN).

**Figure S2.**
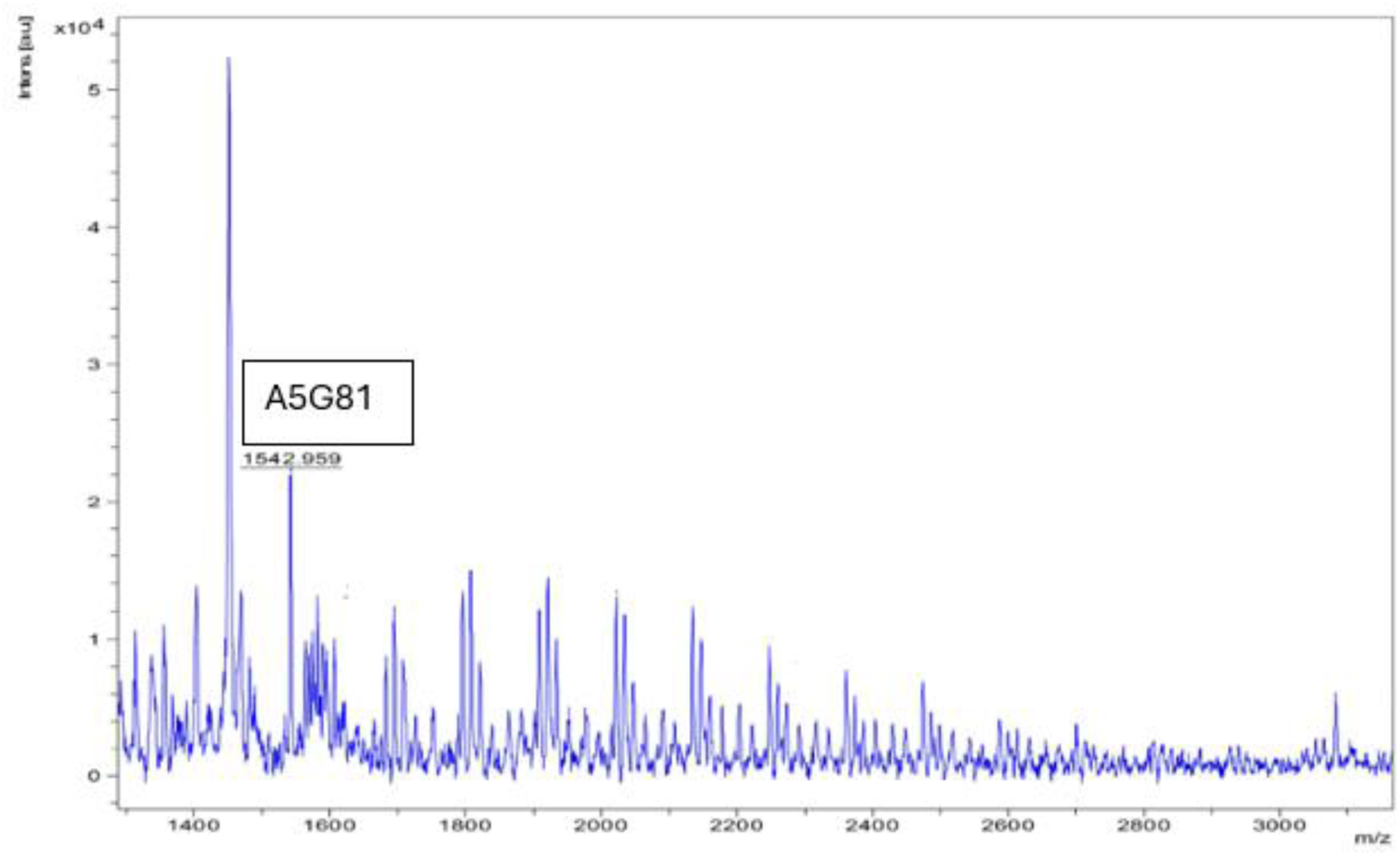
MALDI-TOF spectrum of A5G81-PPCN. Highlighted peaks correspond to A5G81-conjugated species, confirming covalent attachment of the peptide to the PPCN backbone.

**Figure S3.**
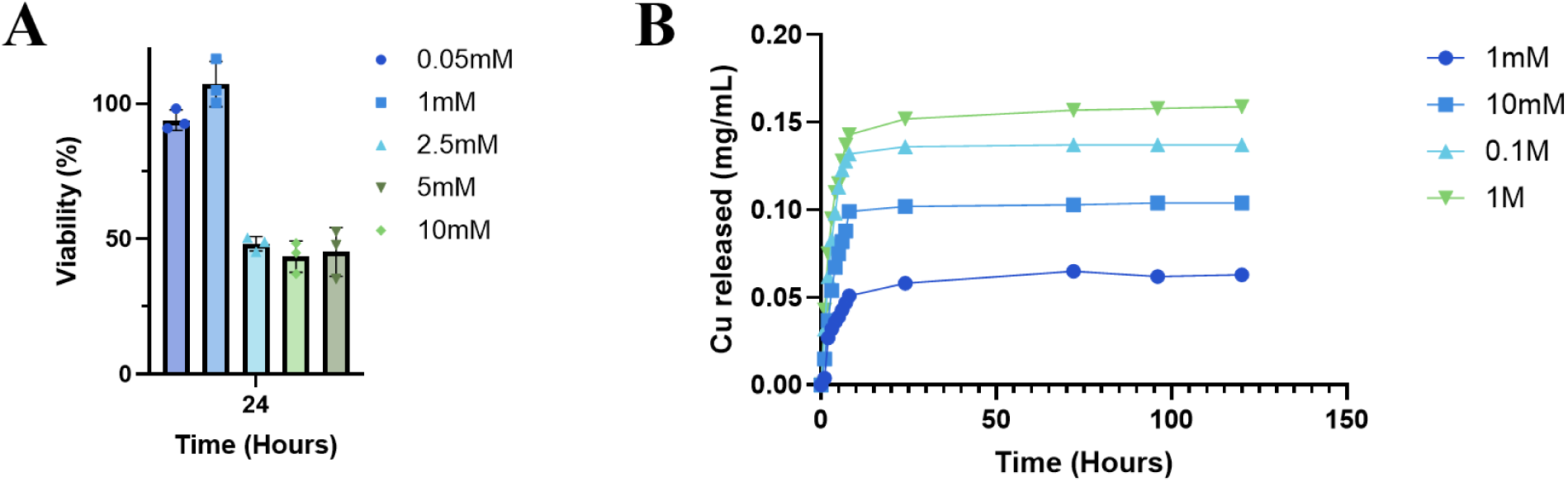
Cell viability and copper release of PPCN-CMOF. (A) Cell viability following 24-hour exposure to varying concentrations of PPCN-CMOF. Viability remained high at lower concentrations, with a notable inverse correlation observed beyond 1 mM copper concentration. (B) Copper release of CMOF from PPCN in PBS.

**Figure S4.**
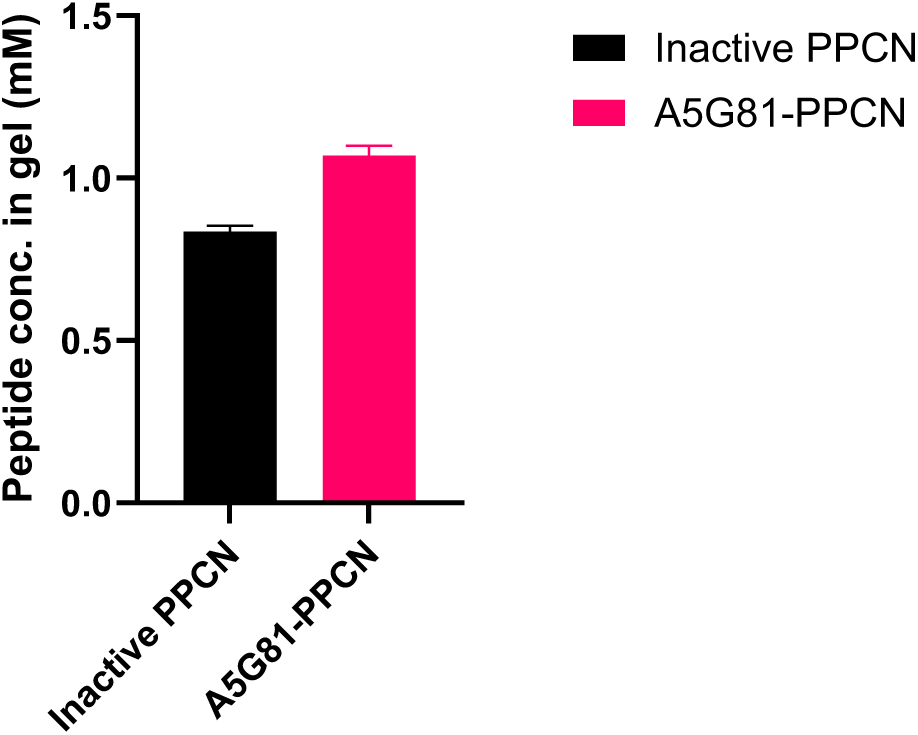
TNBSA assay for A5G81-PPCN. The final peptide concentration in the solution, which was approximately 1 mM.

---

**Table S1.**
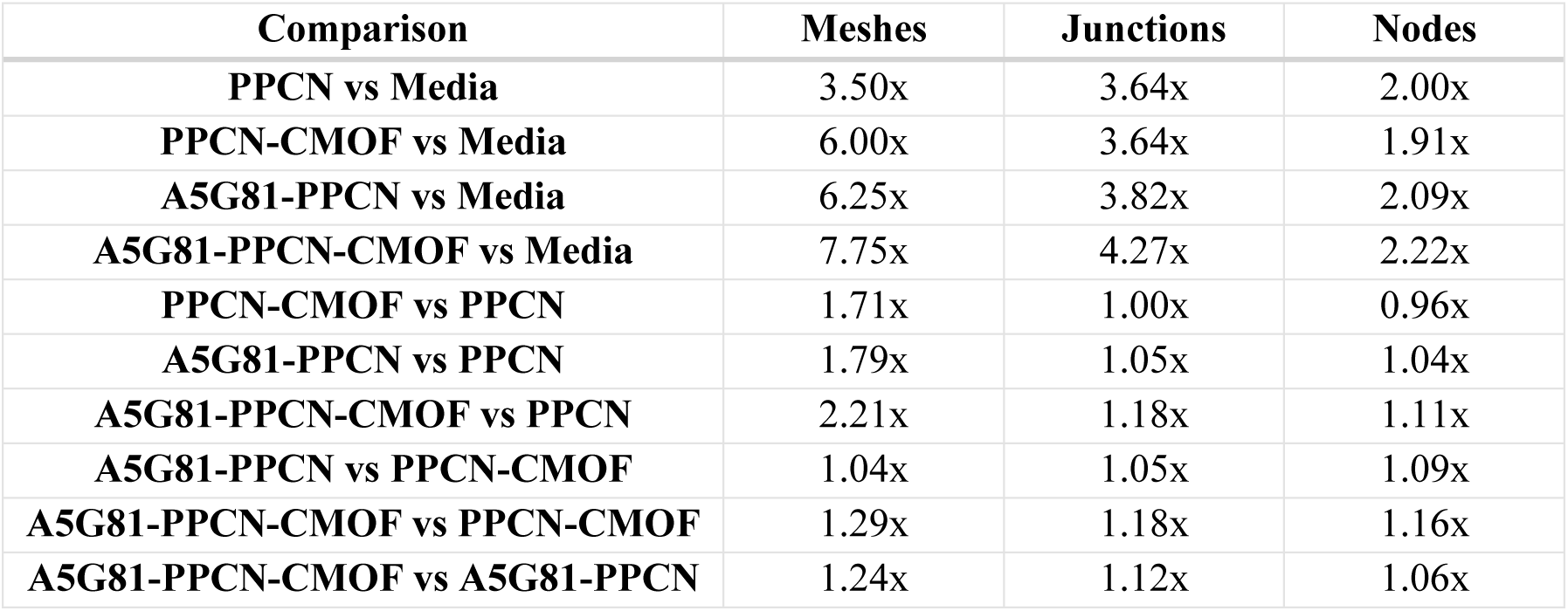
Fold change in tubulogenesis metrics between treatment groups. All four PPCN formulations increased every metric relative to media, by 3.5–7.8× for meshes, 3.6–4.3× for junctions and 1.9–2.2× for nodes. Among formulations, the differences were mainly in mesh formation. PPCN-CMOF, A5G81-PPCN and A5G81-PPCN-CMOF formed 1.7–2.2× more meshes than PPCN alone, and A5G81-PPCN-CMOF gave the highest mesh count. Junctions and nodes differed by no more than ∼1.2× among formulations.

**Table S2.**
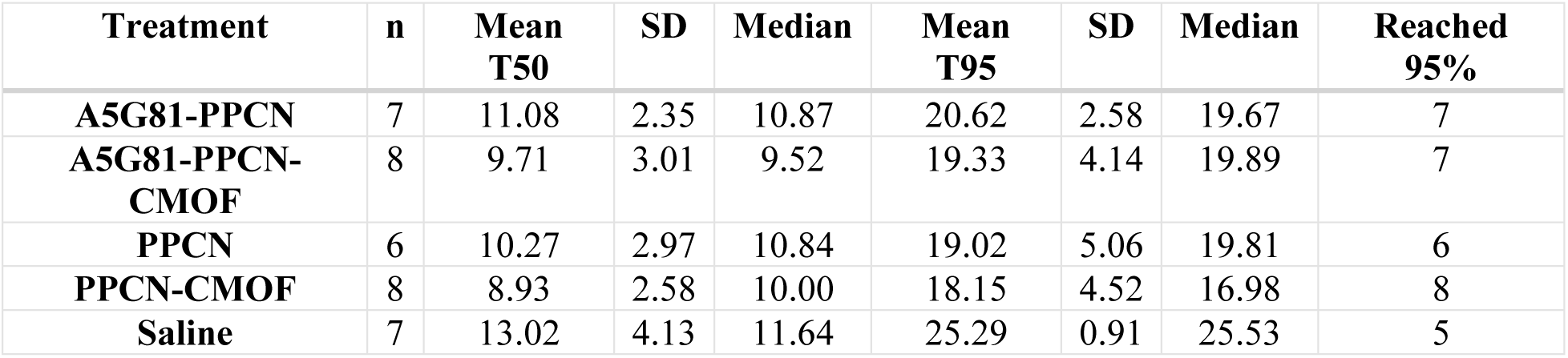
Time to 50% (T50) and 95% (T95) wound closure for non-infected group. T50 and T95 are the time, in days, for the wound area to close by 50% and 95% of its initial area. The time required to achieve 50% (T50) and 95% (T95) wound closure was estimated for each animal using linear interpolation between consecutive wound measurements. Wounds treated with saline, PPCN, PPCN-CMOF, A5G81-PPCN or A5G81-PPCN-CMOF (n = 36 wounds per group). For each threshold, the first interval in which the wound closure percentage crossed the predefined threshold was identified, and the crossing time was estimated assuming a linear change between observations. Animals that did not reach the specified threshold during follow-up were assigned a missing value for that endpoint. T50 and T95 were summarized by treatment group using the mean, standard deviation (SD), and median, and the number of animals reaching 95% wound closure was also reported.

| **Treatment** | **n** | **Mean T50** | **SD** | **Median** | **Mean T95** | **SD** | **Median** | **Reached 95%** |
|---|---|---|---|---|---|---|---|---|
| **A5G81-PPCN** | 7 | 11.08 | 2.35 | 10.87 | 20.62 | 2.58 | 19.67 | 7 |
| **A5G81-PPCN-CMOF** | 8 | 9.71 | 3.01 | 9.52 | 19.33 | 4.14 | 19.89 | 7 |
| **PPCN** | 6 | 10.27 | 2.97 | 10.84 | 19.02 | 5.06 | 19.81 | 6 |
| **PPCN-CMOF** | 8 | 8.93 | 2.58 | 10.00 | 18.15 | 4.52 | 16.98 | 8 |
| **Saline** | 7 | 13.02 | 4.13 | 11.64 | 25.29 | 0.91 | 25.53 | 5 |

**Table S3.**
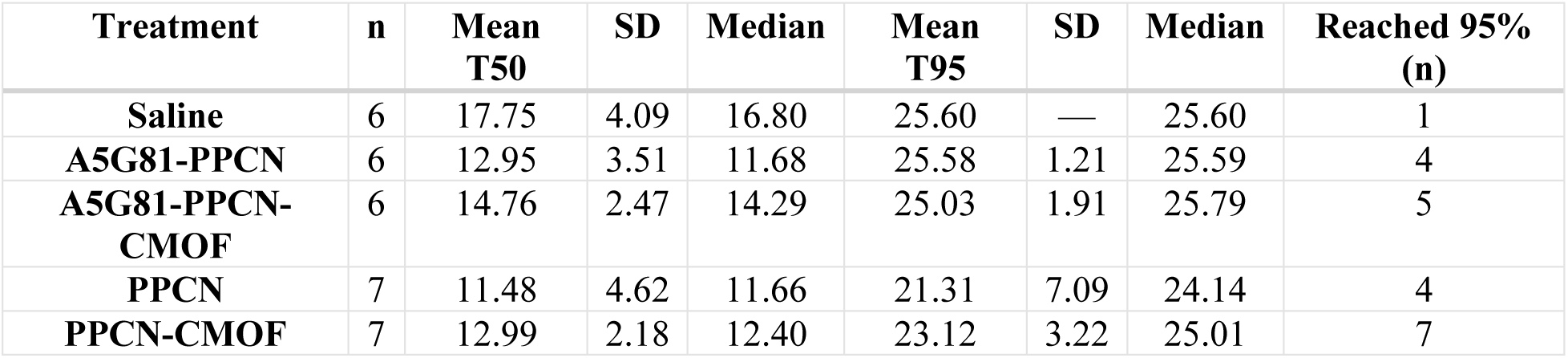
Time to 50% (T50) and 95% (T95) wound closure for infected group. T50 and T95 are the time, in days, for the wound area to close by 50% and 95% of its initial area. The time required to achieve 50% (T50) and 95% (T95) wound closure was estimated for each animal using linear interpolation between consecutive wound measurements. Wounds treated with saline, PPCN, PPCN-CMOF, A5G81-PPCN or A5G81-PPCN-CMOF (n = 32 wounds per group). For each threshold, the first interval in which the wound closure percentage crossed the predefined threshold was identified, and the crossing time was estimated assuming a linear change between observations. Animals that did not reach the specified threshold during follow-up were assigned a missing value for that endpoint. T50 and T95 were summarized by treatment group using the mean, standard deviation (SD), and median, and the number of animals reaching 95% wound closure was also reported.

